# Btk SH2-kinase interface is critical for allosteric kinase activation and its targeting inhibits B-cell neoplasms

**DOI:** 10.1101/862276

**Authors:** Daniel P. Duarte, Allan J. Lamontanara, Giuseppina La Sala, Sukyo Jeong, Yoo-Kyoung Sohn, Alejandro Panjkovich, Sandrine Georgeon, Tim Kükenshöner, Maria J. Marcaida, Florence Pojer, Marco De Vivo, Dmitri Svergun, Hak-Sung Kim, Matteo Dal Peraro, Oliver Hantschel

## Abstract

Bruton’s tyrosine kinase (Btk) is a key component for B-cell maturation and activation. Btk loss-of-function mutations cause human X-linked agammaglobulinemia (XLA). In contrast, constitutive Btk signaling drives several B-cell neoplasms, which may be treated with tyrosine kinase inhibitors (TKIs). Here, we uncovered the molecular mechanism by which a subset of XLA mutations in the SH2 domain strongly perturbs Btk activation. Using a combination of molecular dynamics (MD) simulations and small-angle X-ray scattering (SAXS), we discovered an allosteric interface between the SH2 and kinase domain to which multiple XLA mutations map and which is required for Btk activation. As allosteric interactions provide unique targeting opportunities, we developed an engineered repebody protein binding to the Btk SH2 domain and able to disrupt the SH2-kinase interaction. The repebody prevented activation of wild-type and TKI-resistant Btk, inhibited Btk-dependent signaling and proliferation of malignant B-cells. Therefore, the SH2-kinase interface is critical for Btk activation and a targetable site for allosteric inhibition.

## INTRODUCTION

Almost 2% of the human genes encode for 518 protein kinases (Manning et al., 2002). Due to their central role in normal cellular physiology, the enzymatic activity of kinases is tightly regulated. In contrast, most cancers carry driver or passenger mutations in kinases that cause their aberrant activation and/or display a functional ‘addiction’ to certain kinase pathways for their proliferation/survival. Therefore, kinases are major drug targets and since the clinical approval of imatinib (Gleevec) in 2001 - the first orally available drug inhibiting the BCR-ABL fusion kinase in chronic myelogenous leukemia patients - 47 additional kinase inhibitors were approved (Druker et al., 2001, Hantschel, 2015). A major setback in targeted kinase inhibitor therapy is the development of drug resistance, commonly due to point mutations in the targeted kinase, but also by various other mechanisms (Konieczkowski et al., 2018). For cancers driven by BCR-ABL or EGFR, second-generation drugs were developed that target disease clones carrying resistance mutations, although further resistance by compound mutations or activation of alternative pathways may occur (O’Hare et al., 2012, Jia et al., 2016). An attractive alternative is the targeting of allosteric sites in kinases other than the ATP-binding pocket. These allosteric sites must be critical for the regulation of kinase activity or substrate recruitment to be viable drug targets. Targetable allosteric regulatory sites have been identified for a few kinases and include the myristoyl binding pocket and SH2-kinase interface in BCR-ABL, as well as the PIF pocket in different AGC kinases (Hantschel et al., 2003, Wylie et al., 2017, Grebien et al., 2011, Leroux et al., 2018). As allosteric regulatory pockets are unique to a single kinase or a small class of kinases, one should be able to inhibit oncogenic signaling more selectively and enable either the combinatorial or sequential use of allosteric and ATP-competitive inhibitors, which may diminish or even abolish the outgrowth of resistant clones (Fang et al., 2013, Wylie et al., 2017).

Bruton’s tyrosine kinase (Btk) is a central kinase in B-cell receptor (BCR) signaling that is expressed in the B-cell lineage and in myeloid cells. Loss-of-function mutations in Btk are found in humans with X-linked agammaglobulinemia (XLA). These patients are severely immunocompromised due to the impaired development of B-cells (Vihinen et al., 2000). In contrast, elegant functional genomics work has demonstrated that Btk signaling is critical for the survival of the activated B-cell-like (ABC) subtype of diffuse large B-cell lymphoma (DLBCL) and several other B-cell cancers (Davis et al., 2010, Young and Staudt, 2013). Initial proof-of-concept inhibition of Btk using the FDA-approved BCR-ABL inhibitor dasatinib, which has Btk as one of its major off-targets, triggered the development of more selective Btk inhibitors (Davis et al., 2010, Hantschel et al., 2007, Young and Staudt, 2013). Among those, the first-in-class Btk inhibitor ibrutinib was approved for the treatment of chronic lymphocytic leukemia (CLL), mantle cell lymphoma (MCL), Waldenström’s macroglobulinemia and graft versus host disease since 2013, whereas the more selective drug acalabrutinib was approved in 2017 for MCL. Both drugs suffer from the frequent development of resistance, which is commonly caused by Btk mutations of Cys-481 to which both inhibitors covalently bind, or more rarely by mutations in PLCγ2, downstream of Btk (Quinquenel et al., 2019). Therefore, allosteric mechanisms that regulate Btk activity are particularly attractive as additional drug targets to cope with drug resistance in Btk-dependent B-cell malignancies. Btk and its paralogues Tec, Itk, Bmx and Txk share a conserved SH3-SH2-kinase domain unit with the Src and Abl kinase family of cytoplasmic tyrosine kinases (Hantschel and Superti-Furga, 2004, Shah et al., 2018). Structural and biochemical data showed that intramolecular interactions of the SH3 and SH2 domains with the kinase domain N- and C-lobe, respectively, result in a compact autoinhibited conformation of Btk analogous to Src and Abl kinases (Wang et al., 2015, Shah et al., 2018). In addition, the N-terminal PH-TH domain module of Btk contributes to stabilizing Btk’s autoinhibited conformation (Wang et al., 2015, Joseph et al., 2017). Through inositol phosphate binding to a peripheral site on the PH domain, Btk activation is triggered via dimerization and subsequent trans-autophosphorylation of the kinase domain (Wang et al., 2015). Btk activity is positively regulated by two major phosphorylation events. Tyr-551 in the activation loop can be phosphorylated by upstream Src kinases or trans-autophosphorylated by another Btk molecule. Tyr-223, located in the SH3 domain, is the main Btk autophosphorylation site and thought to be autophosphorylated after Tyr-551 phosphorylation (Park et al., 1996). Although an early small-angle X-ray scattering (SAXS) reconstruction suggested a linear and elongated conformation of active Btk (Marquez et al., 2003), there is little insight on the structural mechanisms and precise molecular events that govern Btk activation.

Here, we show that, based on the analysis of XLA mutations in the SH2 domain, Btk activation critically depends on the formation of an allosteric interface between its SH2 and the N-lobe of kinase domain, which we mapped using a combination of enhanced sampling molecular dynamics (MD) simulations and SAXS. Development of a high-affinity engineered protein antagonist to the Btk SH2 domain targeting its interface with the kinase domain prevents Btk activation in cells, inhibits proliferation and Btk-dependent signaling in malignant B-cells. Therefore, we demonstrate the Btk SH2 domain as alternative allosteric site for therapeutic inhibition of Btk and its most common drug-resistant mutant.

## RESULTS

### A set of XLA mutations preserve canonical Btk SH2 function while impairing kinase activation

Sequencing data indicate that approximately 20% of missense mutations in XLA patients are located within the Btk SH2 domain, but how these mutations result in Btk loss-of-function is poorly understood (Valiaho et al., 2006). While several mutations were shown to decrease protein stability and/or impair canonical phosphotyrosine (pY) peptide binding to the SH2 domain (Mattsson et al., 2000), we were intrigued by a mutational hotspot of surface-exposed residues located on the opposite side of the pY-binding pocket and thus unlikely to be involved in pY-peptide binding (Figure 1A). We first assessed the effects of five representative XLA mutations in this area (K296E, H364D, S371P, R372G, and K374N) and one control XLA mutation (R307G) in the pY binding pocket, on Btk SH2 domain folding, stability, and pY-binding by producing the purified recombinant proteins (Figure 1B, Figure S1A and S1B). Far-UV circular dichroism (CD) spectra and thermal shift analysis demonstrated that the XLA mutations did not significantly change Btk SH2 domain folding and stability compared to the wild-type protein (Figure 1C and Figure S1C). We next determined the effect of the selected XLA mutations on pY-binding in a fluorescence-polarization (FP) binding assay with a labeled pY-peptide. All XLA mutants bound the pY-peptide with similar affinities as the wild-type Btk SH2 domain, whereas the R307G control mutation in the pY binding pocket strongly impaired binding (Figure 1D). Thus, the selected XLA mutations do not act by perturbing folding, stability or pY-binding of the Btk SH2 domain.

**Figure 1.**
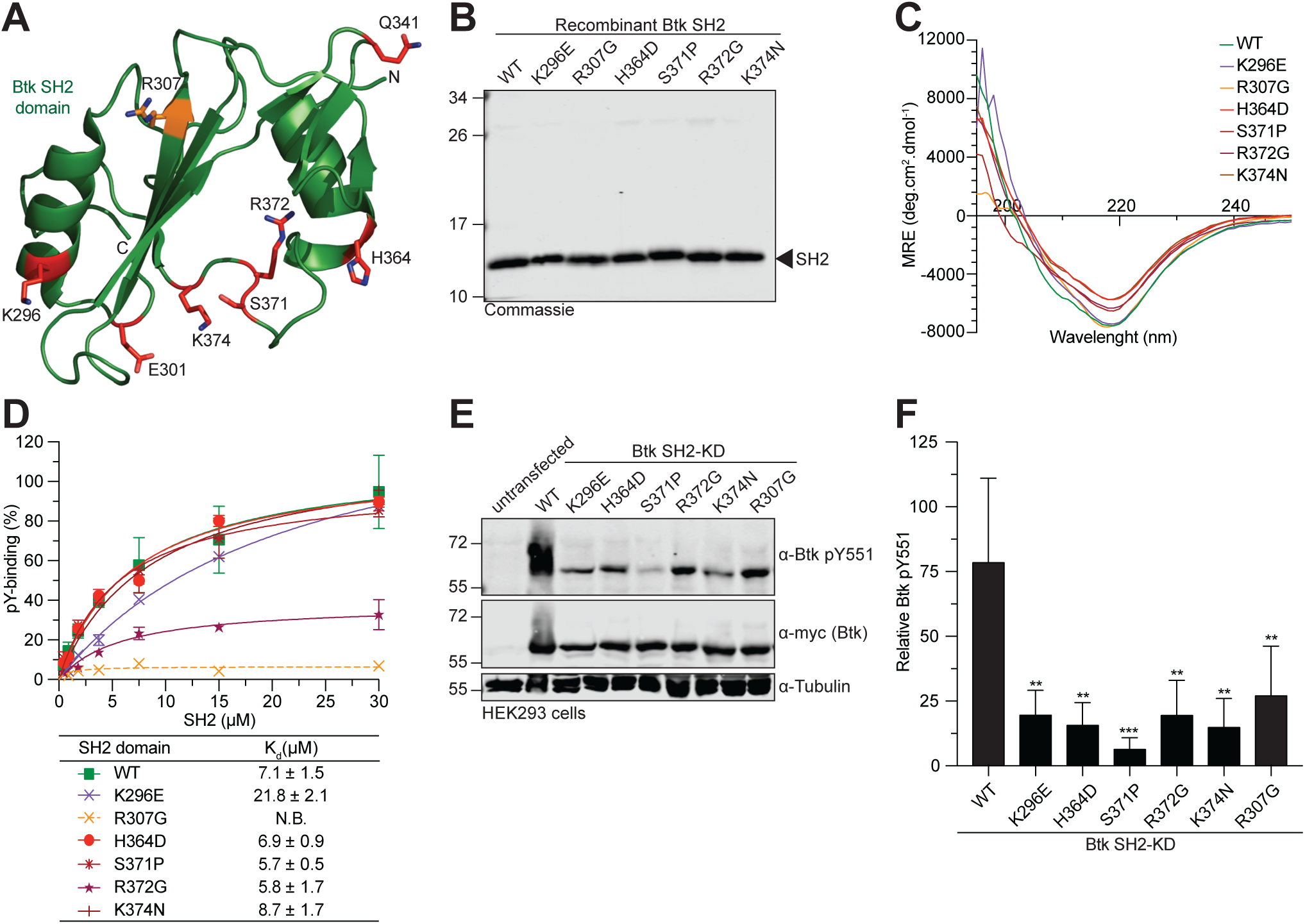
Mutations in Btk SH2 domain abrogates pY551 phosphorylation. A. Mapping of a subset of XLA-patient mutations (red sticks) onto the human Btk SH2 domain structure (PDB 2GE9). The residue R307 (orange sticks) is part of the pY-binding motif (FIV<u>R</u>D). Here and in all subsequent figures, the residue numbering refers to full-length human Btk. B. Representative SDS-PAGE analysis of recombinant wild-type and mutant Btk SH2 domains purified from *E.coli*. C. Averaged far-UV circular dichroism (CD) spectra of recombinant wild-type and XLA mutant Btk SH2 domains. Mean residue ellipticity (MRE) for each protein was calculated from three independent measurements. D. Fluorescence polarization binding assay: Binding of a fluorescently labeled pY-peptide (ADNDpYIIPLPD) to recombinant Btk SH2 domains. Indicated K_d_ values were obtained from at least two technical replicates. The errors indicated are the standard deviations from the curve fitting of the 1:1 binding model. Non-binding (N.B.) E. HEK293 cells were transiently transfected with a construct containing an N-terminal 6xmyc tagged human Btk SH2-KD wild-type or mutants. Immunoblotting of total cell lysates was performed to assess Btk pY551 phosphorylation. F. Quantification of pY551 immunoblot shown in (E) and normalized to total Btk (Myc-Btk) expression. Data shown are the mean ± SD of three biological replicates, and P-values were calculated using an unpaired *t*-test. **P ≤ 0.01 and ***P ≤ 0.001. See also Figure S1.

To determine whether these SH2 mutations affect Btk kinase activity in cells, we introduced these mutations in a Btk SH2-kinase domain (SH2-KD) construct and expressed them in HEK293 cells. Expression of wild-type SH2-KD resulted in robust activation loop phosphorylation (pY551; Figure 1E and 1F). In contrast, all tested XLA mutations strongly decreased phosphorylation at Y551 (Figure 1E and 1F). To test the effect of these mutations on phosphorylation of Y223 within the Btk SH3 domain, which is the most commonly used readout for Btk activity, we introduced them into the larger Btk construct spanning the SH3, SH2 and kinase domains (SH3-SH2-KD). The deleterious effect of an even larger set of XLA mutations in the Btk SH2 domain (Figure S1D) could be corroborated in the SH3-SH2-KD construct with strong impairment of both pY551 and pY223 (Figure S1E and S1F). Importantly, introduction of a control non-XLA mutation (K311E) on the opposite side of the SH2 domain did not affect pY551 and pY223 (Figure S1E and S1F). As expected, when the XLA mutants were introduced in the full-length (autoinhibited) protein, no significant effect on pY551 was observed (Figure S1G).

The intriguing observation that certain XLA mutations do not impact on canonical Btk SH2 domain function suggests the presence of a yet unidentified novel mechanism on how the Btk SH2 domain participates in kinase activation.

### SH2 domain is critical for Btk kinase activation

As the phenotype of the above-described XLA mutations on Btk kinase activation resemble the phenotype of structure-guided targeted mutations in SH2-kinase domain intramolecular interfaces in the Abl and Fes kinases (Filippakopoulos et al., 2008, Grebien et al., 2011), we hypothesized Btk kinase activation by an analogous allosteric mechanism. To address this, we recombinantly expressed sequential domain deletion constructs (Figure 2A) in the presence of YopH phosphatase using baculovirus-infected Sf9 cells to obtain unphosphorylated proteins. All proteins were purified to homogeneity (Figure 2B). Mass spectrometry and immunoblotting analysis confirmed their identity and absence of phosphorylation (Figure S2A and S2B). These recombinant Btk proteins were incubated with Mg^2+^/ATP and *in vitro* autophosphorylation on Y551 was monitored over time (Figure 2C). A kinase-dead Btk SH2-KD protein (D521N) was included as negative control. The SH2-KD construct showed a strong increase in autophosphorylation kinetics compared to the kinase domain alone (KD) and the SH2-KD D521N control (Figure 2D and 2E). In agreement with the crystal structure of mouse Btk SH3-SH2-KD, we observed lower pY551 autophosphorylation than with the SH2-KD construct, but still significantly higher than for the Btk KD, as the presence of the SH3 domain likely induced a more closed autoinhibited conformation of Btk, similar to Abl and Src kinases (Figure 2D and 3E, Wang et al., 2015). The observed lower *in vitro* autophosphorylation for full-length Btk (Figure 2D and 2E) is in line with a recent molecular model of full-length Btk in solution, where the PH-TH domain docks onto the KD to further stabilize its autoinhibition (Joseph et al., 2017). Furthermore, strong total pY phosphorylation of SH2-KD was corroborated in this assay (Figure S2C-E) and corresponds to multiple autophosphorylation sites that we mapped using mass spectrometry (Table S1). Using this assay, we could also show that phosphorylation on Y223 preceded Y551 phosphorylation *in vitro*, which agrees with a previous model where autophosphorylation on Y223 may contribute to full activation of the kinase to further transphosphorylate other Btk molecules on Y551 (Figure S2F; Park et al., 1996).

**Figure 2.**
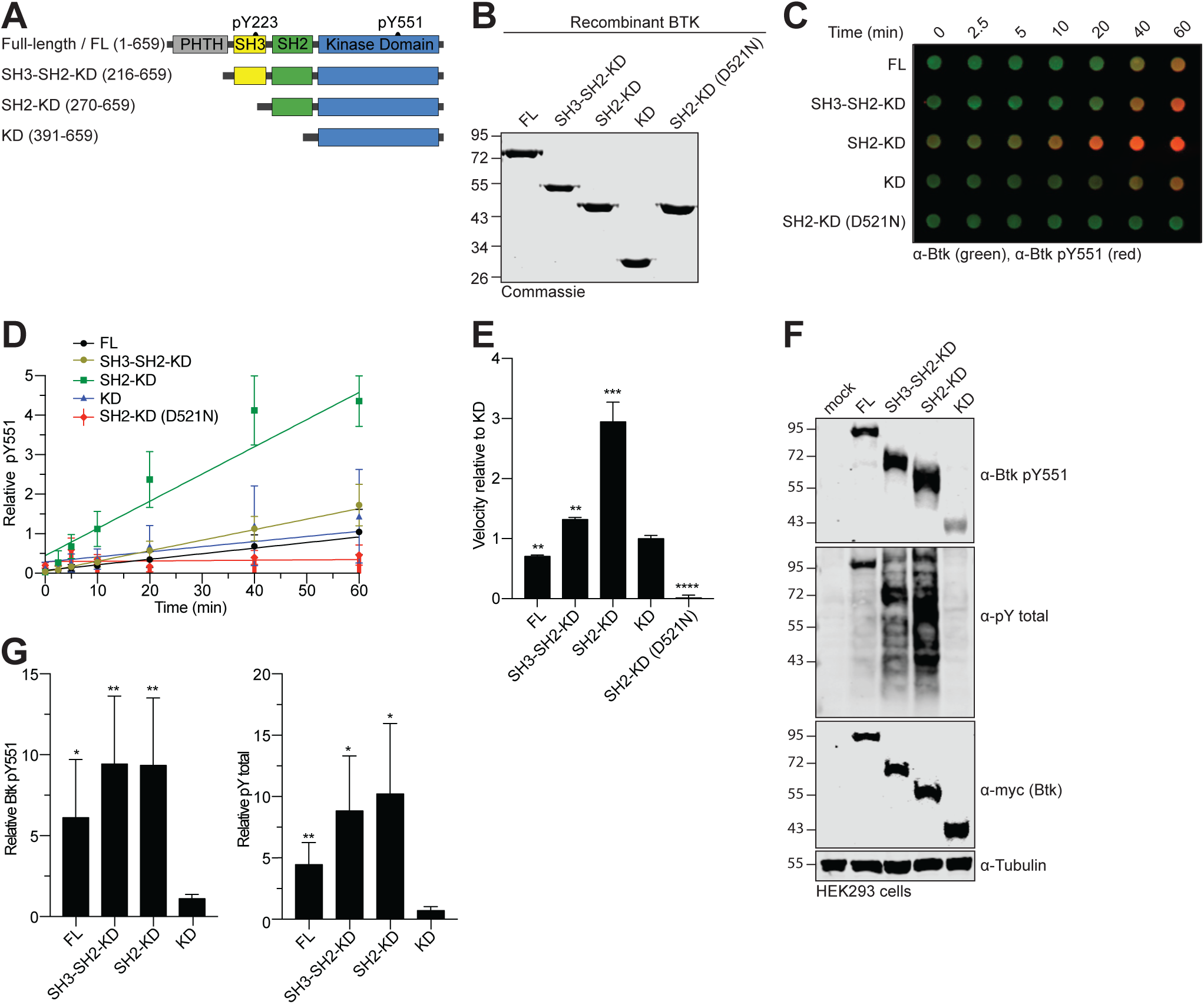
SH2 domain is critical for the activation of Btk kinase. A. Schematic representation of Btk constructs used in this study. Construct/domain boundaries and location of the key activating tyrosine phosphorylation sites (pY223 and pY551) are indicated. B. Representative SDS-PAGE analysis of recombinant untagged Btk proteins purified from Sf9 cells. C. Autophosphorylation assay performed by incubating 1 µM of recombinant Btk proteins with 1 mM ATP and 20 mM Mg^2+^ at room temperature. The levels of pY551 (red channel) and total Btk (green channel) were assessed using immunoblotting in a dot blot apparatus and quantified using the Odyssey Imaging system (Li-Cor). D. Quantitative analysis of Btk autophosphorylation kinetics from dot-blot experiments shown in (B). The ratio of pY551 and total Btk protein is plotted over time. Means ± SD of at least three independent experiments are shown. The slopes (relative velocities) of linear fits were calculated. E. Relative velocities of autophosphorylation relative to the Btk KD are shown. Data are means ± SD of at least three independent experiments. P-values were calculated using an unpaired *t*-test. **P ≤ 0.01, ***P ≤ 0.001 and ****P ≤ 0.0001. F. HEK293 cells were transiently transfected with the indicated Btk constructs containing an N-terminal 6x-Myc tag. Immunoblotting of total cell lysates was performed with the indicated antibodies to assess Btk expression and activation. G. Quantification of pY551 (left) and total pY (right) shown in (E) normalized to total Btk (Myc-Btk) expression and relative to the KD. Data shown are the mean ± SD of three biological replicates. P-values were calculated using an unpaired *t*-test. *P ≤ 0.05, **P ≤ 0.01, ***P ≤ 0.001 and ****P ≤ 0.0001. See also Figure S2.

The strong activating effect of the Btk SH2 domain on autophosphorylation *in vitro* could be corroborated when expressing these constructs in HEK293 cells. Here, the presence of SH2 domain strongly increased Btk autophosphorylation, when compared to KD alone, which shows very low pY551 levels (Figure 2F and 2G). Noteworthy, the SH3-SH2-KD construct showed even higher activation when expressed in cells (Figure 2F and 2G), which could be due to binding of cellular SH3-SH2 ligands that destabilize the autoinhibited conformation of SH3-SH2-KD. This data indicated that the presence of the Btk SH2 domain is critical for the activation of Btk *in vitro* and in cells.

### Btk SH2-KD adopts an elongated conformation to trigger kinase activation

We next turned our focus to investigate the structural basis for SH2-dependent allosteric activation of Btk. In contrast to the Fes, Abl and Csk kinases, where SH2-KD units resembling active conformations could be crystallized and had revealed diverse intramolecular interfaces with the kinase domain N-lobe, we and others have failed to crystallize active Btk. To provide molecular models of the Btk SH2-KD unit, we used enhanced sampling molecular dynamics (MD) simulations. In order to probe for the interaction of the Btk SH2 domain with the KD, we ran multiple replicas of scaled MD simulations for a total of ∼4 µs long trajectories. Scaled MD is an enhanced sampling MD simulation scheme that allows the sampling of µs-ms time-scale events, such as domain-domain binding (Mollica et al., 2015a). This time frame is prohibitive using classical approaches, such as equilibrium MD simulations. Using this approach, we could sample the binding of SH2 to KD in ∼100 ns of simulated time, thus collecting multiple binding events and associated statistics.

Our MD data demonstrated that the SH2 may interact with the KD at different positions, notably at the back, top, and front of the KD N-lobe (Figure S3A). The most representative clusters were located in the back of the KD, followed by a more elongated conformation with the SH2 placed on top of the N-lobe (Fig. 3A). Strikingly, the elongated models suggest that several of the SH2 residues mutated in XLA participate in the interaction interface with the N-lobe of the KD (Figure 3B). Noteworthy, the SH2-KD linker seems to be critical for the interaction with the KD, as our MD simulation without SH2-KD linker failed to probe any SH2-KD interactions (data not shown).

**Figure 3.**
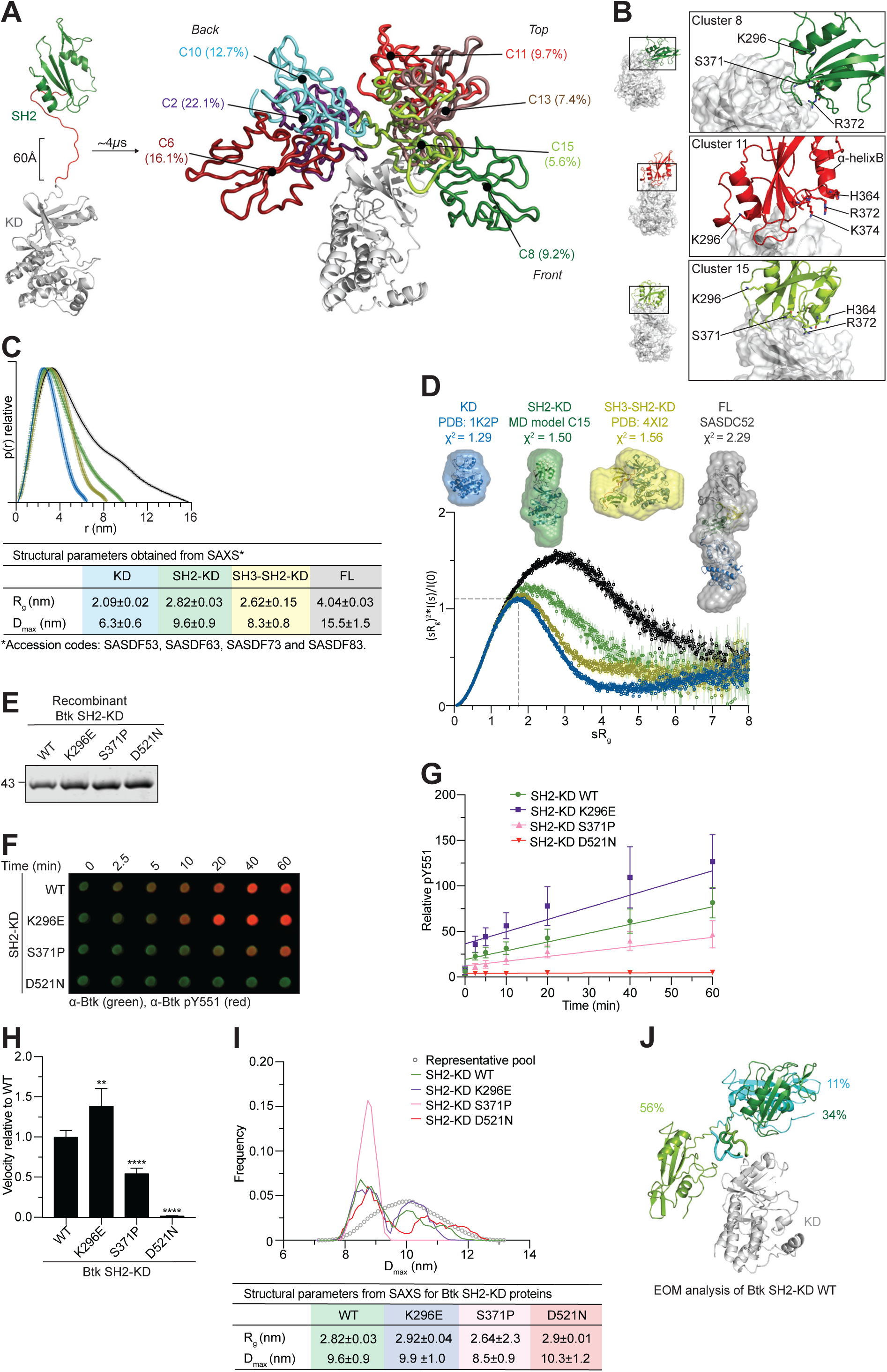
SH2 domain interaction with N-lobe of the KD activates the Btk kinase. A. Scaled MD simulations were performed using a model including the Btk SH2 and KD crystal structures (PDB 6HTF and 1K2P, respectively). The most populated clusters of the SH2 positions (several colors) relative to the KD (white) are shown. The percentages indicate the population of the cluster with respect to the entire simulation time. B. Detailed view of the SH2-KD interface of clusters 8, 11 and 15. SH2 residues mutated in XLA are indicated as sticks. C. Maximal particle dimension (D_max_) of Btk proteins (top) and summary of structural paraments obtained from SAXS (bottom). The table summarizes the particle dimensions (R_g_ and D_max_) and the ± error for the indicated constructs. See Table S2 for details. D. Dimensionless Kratky plot of Btk proteins. The grey dashed line represents the theoretical peak assuming an ideal Guinier region for a globular particle. *Ab initio* envelope reconstructions obtained from SAXS (surface representation) were superimposed on the crystal structures for Btk KD and SH3-SH2-KD (PDB 1K2P and 4XI2, respectively) are shown on top of the Kratky plot. For the SH2-KD protein, the structure of an elongated MD model with the best agreement with the experimental SAXS data is shown (model C15, see panels (A) and (B)). FL protein shows an extended conformation as observed in a previously published SAXS reconstruction (SASDC52). E. Representative SDS-PAGE analysis of recombinant untagged SH2-KD proteins purified from Sf9 cells. F. Autophosphorylation assay performed by incubating 1 µM and 20 mM Mg^2+^ of recombinant Btk proteins with 1 mM ATP at room temperature. The levels of pY551 (red channel) and total Btk (green channel) were assessed using immunoblotting in a dot blot apparatus and quantified using the Odyssey Imaging system (Li-Cor). G. Quantitative analysis of Btk autophosphorylation kinetics from dot-blot experiments shown in (F). The ratio of pY551 and total Btk protein is plotted over time. Means ± SD of three independent experiments are shown. The slopes (relative velocities) of linear fits were calculated. H. Relative velocities of autophosphorylation relative to the Btk KD are shown. Data show the mean ± SD of three independent experiments. P values relative to wild-type were calculated using an unpaired *t*-test. **P ≤ 0.01, and ****P ≤ 0.0001. I. Flexibility analysis of SH2-KD based on ensemble of optimization method (EOM 2.0) using the experimental SAXS data for mutant Btk proteins. On top, the maximal particle dimension (D_max_) of selected conformers for each protein (lines) from a representative pool of theoretical conformations (dotted line) is shown. Below, a table shows the summary of the structural parameters obtained for wildtype and mutant proteins. J. Structural representation of selected conformers of SH2-KD wild-type protein based on the EOM 2.0 analysis shown in (I) (green line). The percentages represent the contribution of each conformer to a SAXS profile in good agreement with the experimental curve. See also Figure S3 and Table S2.

In order to provide an unbiased and independent experimental validation of the MD model, we performed an extensive analysis of multiple recombinant Btk proteins with small-angle X-ray scattering (SAXS). This method allows to directly reconstruct low-resolution particle shapes *ab initio*, and also to study conformational dynamics of multidomain proteins and complexes in solution. From the SAXS data, the SH2-KD construct adopts a more elongated conformation with increased particle dimension (D_max_) in comparison to the KD and SH3-SH2-KD proteins (Figure 3C and Figure S3B). The extended conformation of the SH2-KD protein is independent of kinase activity, as also seen with a kinase-inactive mutant (D521N; Table S2), and consistently revealed by batch and also size exclusion chromatography coupled to SAXS (SEC-SAXS) measurements (data not shown). In addition to the increased particle dimension, the normalized Kratky plot suggests that SH2-KD is flexible compared to the globular KD and SH3-SH2-KD proteins (Figure 3D, bottom). *Ab initio* shape reconstructions from SAXS show an excellent agreement with the available KD and (closed autoinhibited) SH3-SH2-KD crystal structures (PDB 1K2P and 4XI2, respectively, Figure 3D, top). Importantly, the *ab initio* envelopes of SH2-KD can be superimposed very well with the elongated MD models (e.g. C15, χ^2^=1.50) in which the SH2 domain is interacting with the N-lobe of the KD (Figure 3D). The full-length protein has been previously reported to adopt an equilibrium of conformations with a predominant compact and autoinhibited state in solution (Joseph et al., 2017), whereas our SAXS data from the full-length protein is compatible with its extended conformation and agrees with a previous report (Marquez et al., 2003). Overall, independent use of MD and SAXS supports an extended model for the Btk SH2-KD, where the SH2 is placed on top and interacts predominantly with the N-lobe of the KD. To further probe the validity of this model, we took advantage of XLA mutations at different positions on the SH2 surface. Based on our SH2-KD model, S371 is part of the interface with the KD N-lobe, while K296 is solvent-exposed and does not participate in this interdomain interaction (model C15, Figure 3B). When introduced into Btk SH2-KD, the recombinant purified S371P protein showed decreased *in vitro* autophosphorylation kinetic on Y551 compared to the wild-type protein (Figure 3E-H). Interestingly, the K296E mutant protein showed a mild increase in Y551 autophosphorylation compared to wild-type, as the mutation may disfavor a more inactive conformation of the SH2-KD protein (Figure 3E-H). Total pY phosphorylation of the S371P protein was also impaired and further supports the lower autophosphorylation capacity observed for this construct (Figure S3D-G). To provide additional insights on how these mutations affect SH2-KD conformation, we assessed protein flexibility using SAXS data combined with the ensemble optimization method (EOM; Tria et al., 2013). EOM generates a large number of conformations (>10,000) using the KD and SH2 domain structures taking the native linker into account, calculates theoretical SAXS curves for all models, and selects a mixture of conformers that fits the experimental SAXS data. First, SAXS data from the mutants indicated that the S371P protein is somewhat more compact than wild-type, K296E and D521N proteins (Figure S3H and S3I, Table S2). EOM analysis revealed that SH2-KD wild-type is rather flexible with few different conformers co-existing in solution, but indicating extended conformations with the SH2 placed on top of the kinase (Figure 3I and 3J). Analysis of the kinase-dead D521N and K296E SH2-KD proteins indicate similar flexibility as for the wild-type protein and also similar overall molecular dimensions (R_g_ and D_max_). In contrast, the S371P mutant protein showed a shift towards more compact conformation, with a decrease in R_g_ and D_max_ compared to the wild-type and mutations not affecting the interdomain interface (Figure 3I). Importantly, measurements of the SH2-KD S371P protein using the purified protein in batch mode as well as SEC-SAXS showed similar features (data not shown). The compact conformation adopted by the SH2-KD S371P is consistent with the decreased phosphorylation observed in the autophosphorylation assay (Figure 3E-H). Noteworthy, EOM analysis performed for the SH3-SH2-KD and full-length wild-type proteins is consistent with the models showing, respectively, a compact and a mixture of compact and elongated conformations in solution (Figure S3K). Summarizing, we provide a model for Btk activation via allosteric interaction of the SH2 domain predominantly placed on top of KD. Although the SH2-KD interaction seems less sturdy as in Abl and Fes, a molecular model for the interface of SH2 interacting with the KD can be deduced from our data.

### Development of a protein binder targeting the BTK SH2 domain

In order to demonstrate the importance of the proposed allosteric interaction of the Btk SH2 domain with the kinase domain in regulating Btk activity, we developed a repebody binder. Repebodies are engineered non-antibody scaffold proteins composed of leucine-rich repeat (LRR) modules that can be engineered to bind targets with high specificity. A human Btk SH2 domain-targeting repebody, termed rF10, was generated using phage display selection and affinity maturation (Lee et al., 2012; Figure S4A). rF10 and a non-binding control repebody (rNB) readily purified from *E.coli* (Figure 4A). The affinity of rF10 to the Btk SH2 domain is ∼15 nM with a binding stoichiometry of 1:1 (Figure 4B). In contrast, rF10 showed no binding to the SH2 domains from its close relatives, the tyrosine kinases Abl and Lck, demonstrating a >500-fold selectivity for the Btk SH2 domain (Figure 4B). Consistent with a high-affinity interaction, a stable 1:1 rF10-SH2 complex could be recovered by size-exclusion chromatography either in complex with the Btk SH2 domain alone (Figure 4C and 4D), as well as the SH2-KD and full-length Btk proteins (data not shown).

**Figure 4.**
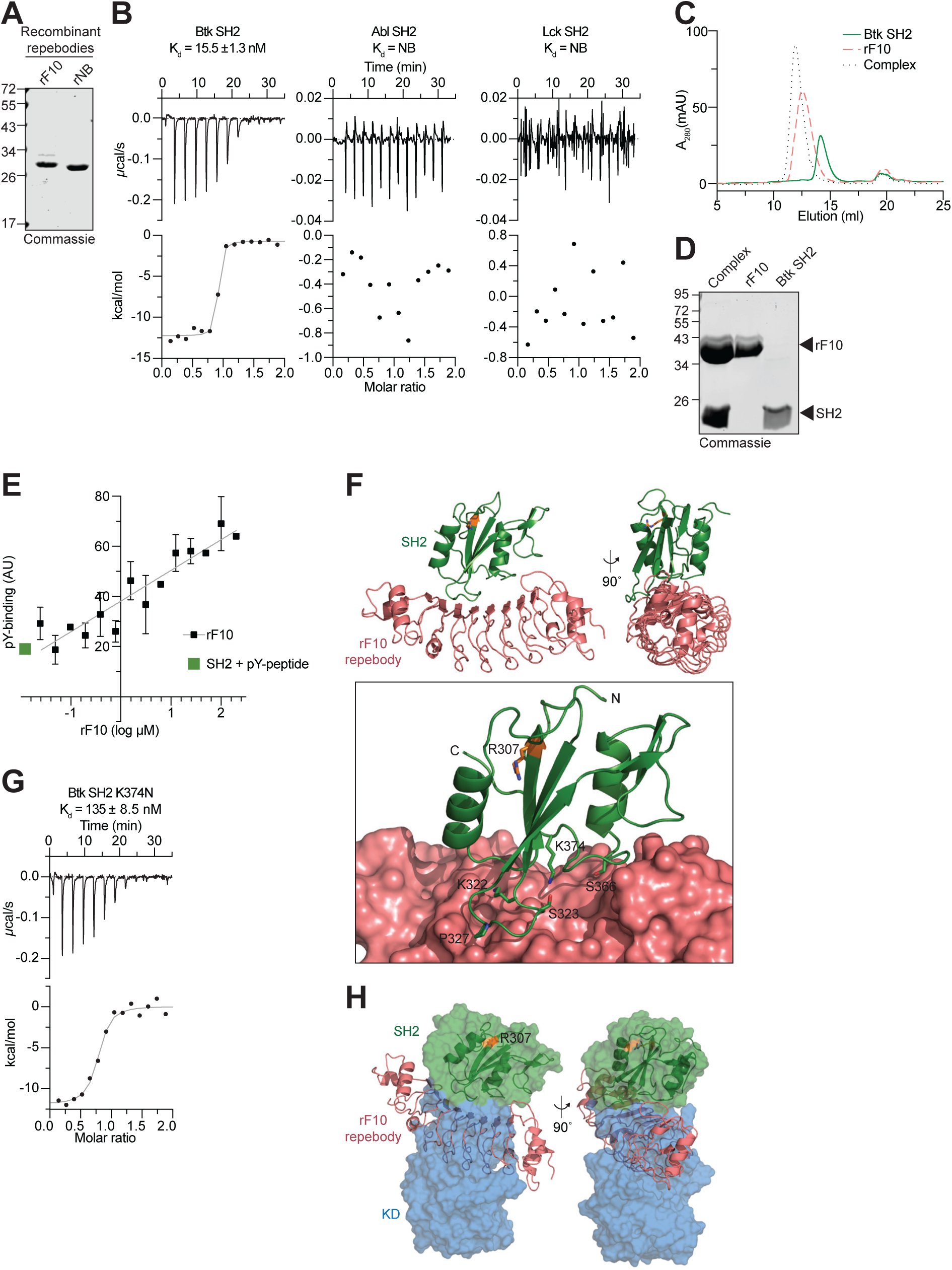
Development of a high-affinity protein binder to the human Btk SH2 domain. A. Representative SDS-PAGE analysis of recombinant repebodies rF10 and rNB purified from *E.coli*. B. ITC measurements of rF10 repebody to the SH2 domains of Btk, Abl and Lck kinases. The top panels show the raw signal from a representative measurement. The bottom panels show the integrated calorimetric data of the area of each peak. The continuous line indicates the best fit to the experimental data assuming a 1:1 binding model. Reported K_d_ value for Btk was calculated from three independent measurements. No binding (NB) K_d_ indicates no interaction between proteins. C. Size-exclusion chromatogram (SEC) analysis of Btk SH2 and rF10 alone, and the SH2-rF10 complex formed by pre-incubation of SH2 and rF10 prior to column injection. D. A fraction of each respective peak from the SEC analysis shown in (C) was resolved by SDS-PAGE and stained with Coomassie. E. Binding-competition assay using fluorescently labeled pY-peptide (ADNDpYIIPLPD) to recombinant Btk SH2 domain in the presence of various concentrations of rF10 repebody. Averages ± SD from three technical replicates are plotted. F. Cartoon representation of the crystal structure of human Btk SH2 (green) in complex with rF10 repebody (salmon), PDB 6HTF. Structural statistics are reported in Table S3. The lower panel shows the rF10 in surface representation, residue R307 (orange) indicates the position of the pY-binding site, and side chains of SH2 residues interacting with rF10 are shown as green sticks. G. ITC measurement of rF10 repebody to Btk SH2 K374N performed as in (A). The K_d_ value was calculated from two independent measurements. H. Superimposition of the rF10-SH2 structure (cartoon representation, color scheme as in (F) on the active Btk SH2-KD model (surface representation, SH2 in green and KD in blue). See also Figure S4 and Table S3.

As other engineered SH2 binders, in particular monobodies (Wojcik et al., 2010, Sha et al., 2013, Kukenshoner et al., 2017), predominantly target the pY peptide binding site, we first tested whether the rF10 repebody interferes with pY-peptide binding using an FP binding assay. Even in the presence of a 20-fold molar excess of rF10, no competition with pY-peptide binding was observed (Figure 4E), indicating that rF10 has a different binding epitope on the SH2 domain. The observed increased FP signal with increasing rF10 concentrations is consistent with the formation of a ternary complex (SH2-pY-peptide-rF10; Figure 4E).

We next solved the crystal structure of the Btk SH2-rF10 complex at 1.9Å resolution (Figure 4F, Table S3, PDB 6HTF), which is the first crystal structure of the isolated human Btk SH2 domain. The overall SH2 domain conformation is very similar to the previously published NMR structure of human Btk SH2 domain (PDB 2GE9, Huang et al., 2006), indicating that rF10 binding does not result in major conformational changes (Wojcik et al., 2016, Kukenshoner et al., 2017). Consistent with the ITC and pY binding assays, the rF10-SH2 crystal structure indicates that rF10 binds to SH2 domain in a 1:1 interaction and the interaction does not involve the pY-binding groove. rF10 binds to multiple residues from the SH2 domain BC loop (K322, S323, G325 and P327) and the C-terminus of the α-helix B (S366 and K374), and it buries a surface area of 2274 Å^2^ (Figure 4F). To further corroborate the rF10 interaction site, a recombinant SH2 containing the XLA mutation K374N in the interface between the SH2-rF10 showed a ∼10-fold decreased affinity (Figure 4G) compared to the wild-type SH2 domain.

Superimposition of the Btk SH2-rF10 structure with a representative SH2-KD structure (MD model C15) revealed dramatic steric clashes of the KD and rF10 (Figure 4H, Video 1). The coincidental strong overlap of the rF10 binding epitope with the proposed Btk SH2-kinase interface led us to hypothesize that rF10 may abrogate the SH2-KD interaction and thereby act as an allosteric Btk antagonist. To probe this hypothesis, we first performed SAXS analysis of rF10 alone and in complex with several Btk constructs (Figure S4B and S4C, Table S4). In line with our hypothesis, SAXS-based reconstructions of rF10-Btk complexes indicate that the conformation of SH2-KD is altered and the interdomain interface is disrupted (Figure S4D). This observation encouraged us to further investigate the functional effects of the rF10 repebody on Btk activity and signaling *in vitro* and in cells.

### Targeting the SH2-KD interface with rF10 results in Btk kinase inhibition

As rF10 was confirmed to bind the Btk SH2 domain, we first measured the binding affinity of rF10 to recombinant SH2-KD, SH3-SH2-KD and full-length Btk. rF10 was found to bind all three proteins with similar low nanomolar affinities (Fig. S5A). We next performed *in vitro* autophosphorylation assays using different recombinant Btk constructs in the presence of a 2-fold molar excess of rF10 or a non-binding repebody (rNB) control (Figure 5A and 5B). rF10 showed a strong inhibitory effect on pY551 autophosphorylation of all tested Btk constructs containing the SH2 domain (full-length, SH3-SH2-KD and SH2-KD; Figure 5C and 5D). Interestingly, even though the constructs SH3-SH2-KD and full-length Btk adopt an autoinhibited conformation with low autophosphorylation activity (see Figure 2C-E), rF10 strongly decreased their remaining activity (Figure 5C and 5D). Consistent with this data also total pY phosphorylation of Btk was decreased in the presence of rF10 (Figure S5B-E).

**Figure 5.**
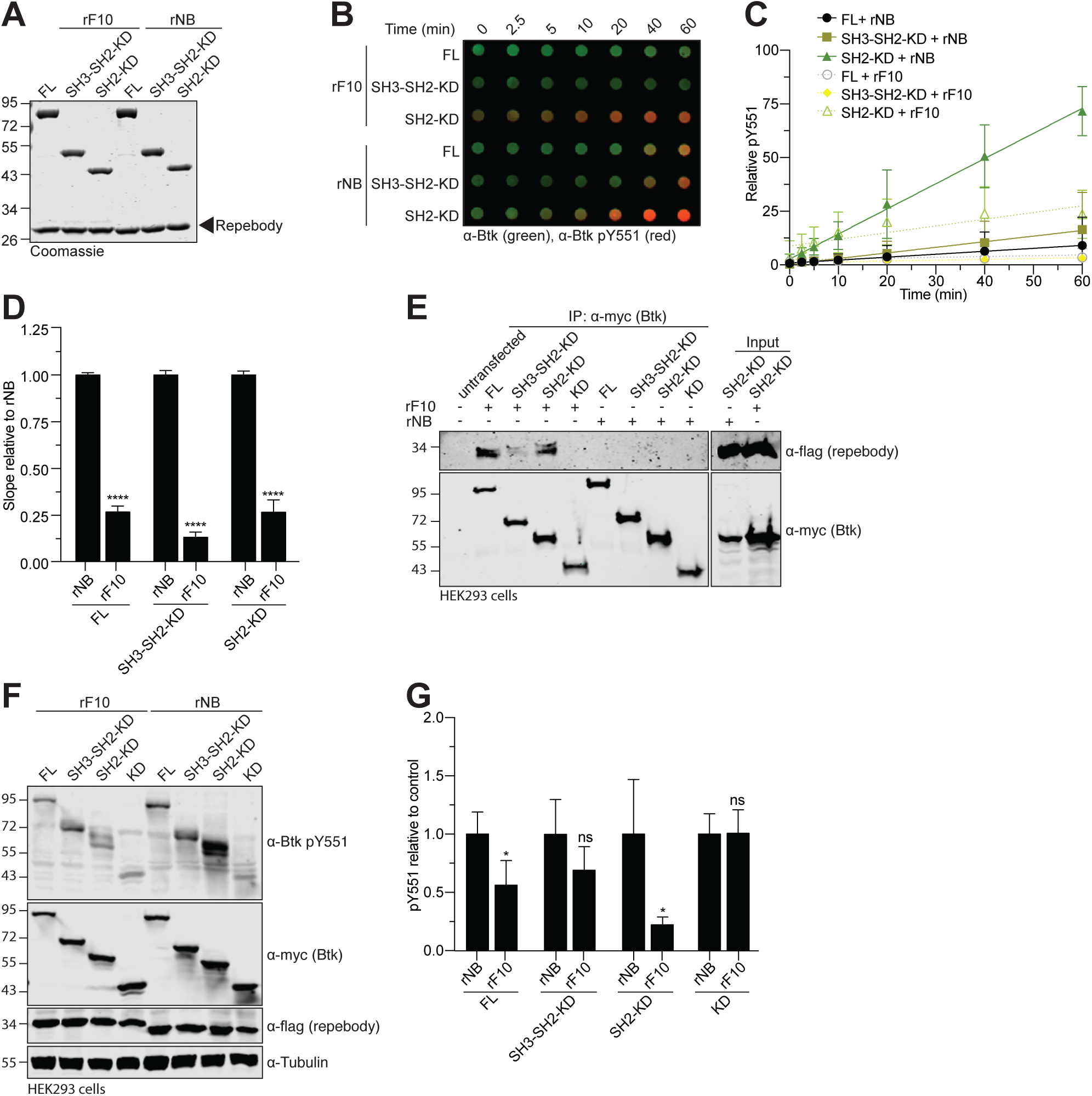
rF10 repebody inhibits Btk activation *in vitro* and in cells. A. Representative SDS-PAGE analysis of recombinant Btk proteins mixed with rF10 and rNB control repebodies and used for *in vitro* autophosphorylation assays. B. Autophosphorylation assay performed by incubation of 1 µM of recombinant human Btk protein and 2 µM of rF10 (dashed lines) or non-binding control repebody (rNB, continuous lines) with 1mM ATP and 20 mM Mg^2+^ at room temperature. The levels of pY551 (red channel) and total Btk (green channel) were assessed using immunoblotting in a dot blot apparatus and quantified using the Odyssey Imaging system (Li-Cor). C. Quantitative analysis of Btk autophosphorylation kinetics in the presence of rF10 (dashed lines) or rNB (continuous lines) repebodies from dot-blot experiments shown in (B). The ratio of pY551 and total Btk protein is plotted over time. Means ± SD of three independent experiments are shown. The slopes (relative velocities) of linear fits were calculated. D. Relative velocities of autophosphorylation for each Btk construct and relative to control repebody are shown. Data show the mean ± SD of three independent experiments. P-values relative to each control rNB repebody were calculated using an unpaired *t*-test. ****P ≤ 0.0001. E. HEK293 cells were transiently co-transfected with indicated Btk constructs and repebodies, and lysates were subjected to immunoprecipitation using anti-Myc coated beads. A representative sample of lysate for each repebody was loaded as expression control. F. Immunoblot analysis of lysates from HEK293 cells transiently co-transfected with indicated Btk constructs and repebodies used to assess Btk pY551 phosphorylation. G. Quantification of pY551 shown in (F) and normalized to total Btk (Myc-Btk) expression level and relative to control repebody. Data shown are the mean ± SD of three biological replicates, and P-values were calculated against each control rNB repebody using unpaired *t*-test. *P ≤ 0.05 and non-significant (ns). See also Figure S5.

To test the ability of rF10 to act an allosteric Btk inhibitor in cells, we first tested whether rF10 interacted with different Btk constructs in mammalian cells. Pull-down assays showed interactions of rF10, but not of the rNB control repebody, with all Btk constructs containing the SH2 domain, but not with the Btk KD (Figure 5E). In the presence of rF10, lower levels of Btk pY551 were observed than when equal amounts of rNB were expressed. Phosphorylation of the KD alone was unaltered in the presence of repebodies, indicating selective SH2 domain-dependent inhibition of Btk autophosphorylation by rF10 (Figure 5F and 5G). To determine the effect of allosteric inhibition of Btk by rF10 on kinase activity, we performed *in vitro* kinase assays with a substrate peptide encompassing Tyr-753 of PLCγ2, a canonical Btk substrate. In the presence of rF10, but not rNB, kinase activity of full-length Btk was strongly inhibited (Figure S5H).

This data collectively showed that targeting the Btk SH2 with a repebody binder at the proposed SH2-kinase interface selectively inhibits Btk activity.

### rF10 selectively decreases viability and inhibits signaling of lymphoma cells

Finally, we investigated whether targeting the SH2-KD interface is sufficient to inhibit Btk activity in neoplastic B-cells. We selected cell lines that express wild-type Btk and are sensitive to ibrutinib (Figure S6A) and transduced them to inducibly express rF10 or the rNB. Upon induction of rF10 expression, we observed a more than 10-fold reduction in cumulative cell numbers as compared to rNB, all uninduced conditions or parental cells (Figure 6A). This was accompanied by a dramatic increase in apoptosis, comparable to the treatment of parental cells with ibrutinib (Figure 6B). Also in TMD8 cells, a decrease of the cumulative cell numbers were observed (Figure S6B).

**Figure 6.**
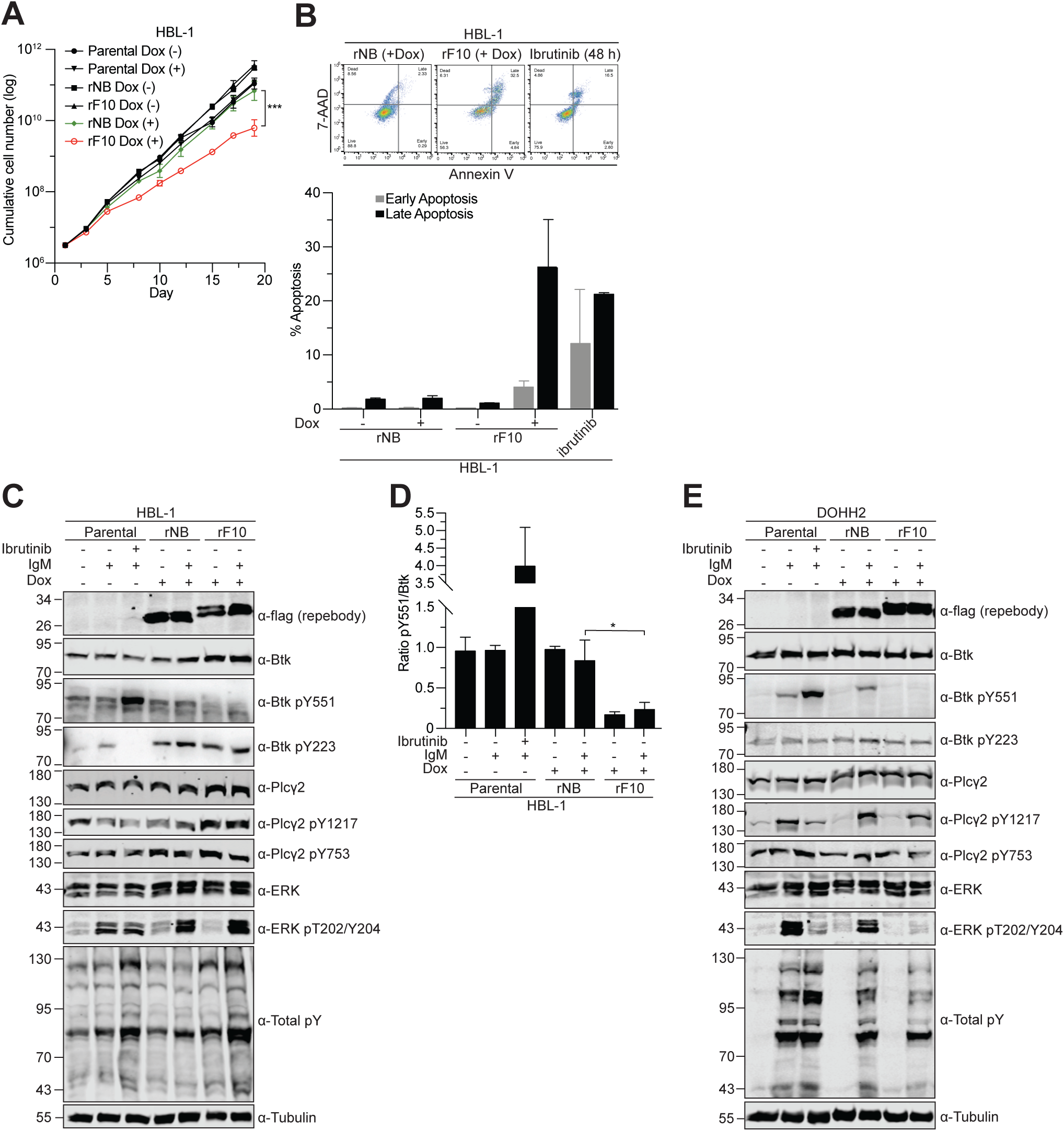
Targeting the Btk SH2-KD interface decreases the viability of B-cell lymphoma cells and inhibits BCR signaling. A. HBL-1 cells were lentivirally transduced with a doxycycline-inducible system for expression of repebodies, and cumulative cell numbers monitored upon treatment with 2 µg mL^-1^ of doxycycline. Parental cells are non-transduced cells. B. HBL-1 inducibly expressing rF10 or rNB control for 7 days were stained with 7AAD and Annexin V to analyze apoptosis by FACS. On top, a representative FACS staining is shown. On the bottom, the quantification of early (7AAD-/Annexin V+) and late (7AAD+/Annexin V+) apoptotic cells obtained from two independent experiments. HBL-1 parental cells treated with 10 µM of ibrutinib for 48 hours were used as positive control. C. Expression of repebodies (flag-tagged) was induced for 48 hours in HBL-1 cells, and BCR signaling stimulated with anti-human IgM or mock-treated for 2 minutes before cell lysis. Ibrutinib treatment (100 nM) was performed for 15 minutes prior to anti-IgM stimulation. Immunoblot analysis of whole-cell lysates with the indicated antibodies is shown. Tubulin was used as loading control. D. Quantification of Btk pY551 shown in (C) and normalized to total Btk expression. Data shown are the mean ± SD from two biological replicates, and P-values were calculated using an unpaired *t*-test. *P ≤ 0.05. E. Expression of repebodies (flag-tagged) was induced for 48 hours in DOHH2 cells, and BCR signaling stimulated with anti-human IgG or mock-treated for 2 minutes before cell lysis. Ibrutinib treatment (100 nM) was performed for 15 minutes prior to anti-IgM stimulation. Immunoblot analysis of whole-cell lysates with the indicated antibodies is shown. Tubulin was used as loading control. See also Figure S6.

rF10 expression promotes decreased Btk pY551 phosphorylation in HBL-1 cells, which could not be recovered by BCR stimulation using anti-human IgM (Figure 6C and 6D). Interestingly, an increase in Btk and PLCγ2 protein levels was observed upon rF10 expression, which may suggest a compensatory mechanism to counteract the activity of rF10 on Btk inhibition (Figure 6C and S6C). The rF10 effects on BCR signaling were also consistent in DOHH2 cells. Here, rF10 expression resulted in decreased Btk pY551, even upon BCR stimulation, decreased PLCγ2 phosphorylation on Y1217, one of the two main Btk phosphorylation sites, as well as strongly decreased in Erk activation (Figure 6E). Despite the strong inhibition of the BCR signaling pathways by rF10, we did not observe growth inhibition of DOHH2 cells when targeting the SH2-KD interface using the rF10 (data not shown).

We next tested whether targeting the SH2-KD interface offers an alternative approach to target TKI-resistant Btk. Importantly, rF10 was able to decrease pY551 (Figure 7A and 7B) and total pY (Figure S7A and S7B) of different Btk C481S constructs expressed in HEK293 cells and to a similar extent than of wild-type Btk.

**Figure 7.**
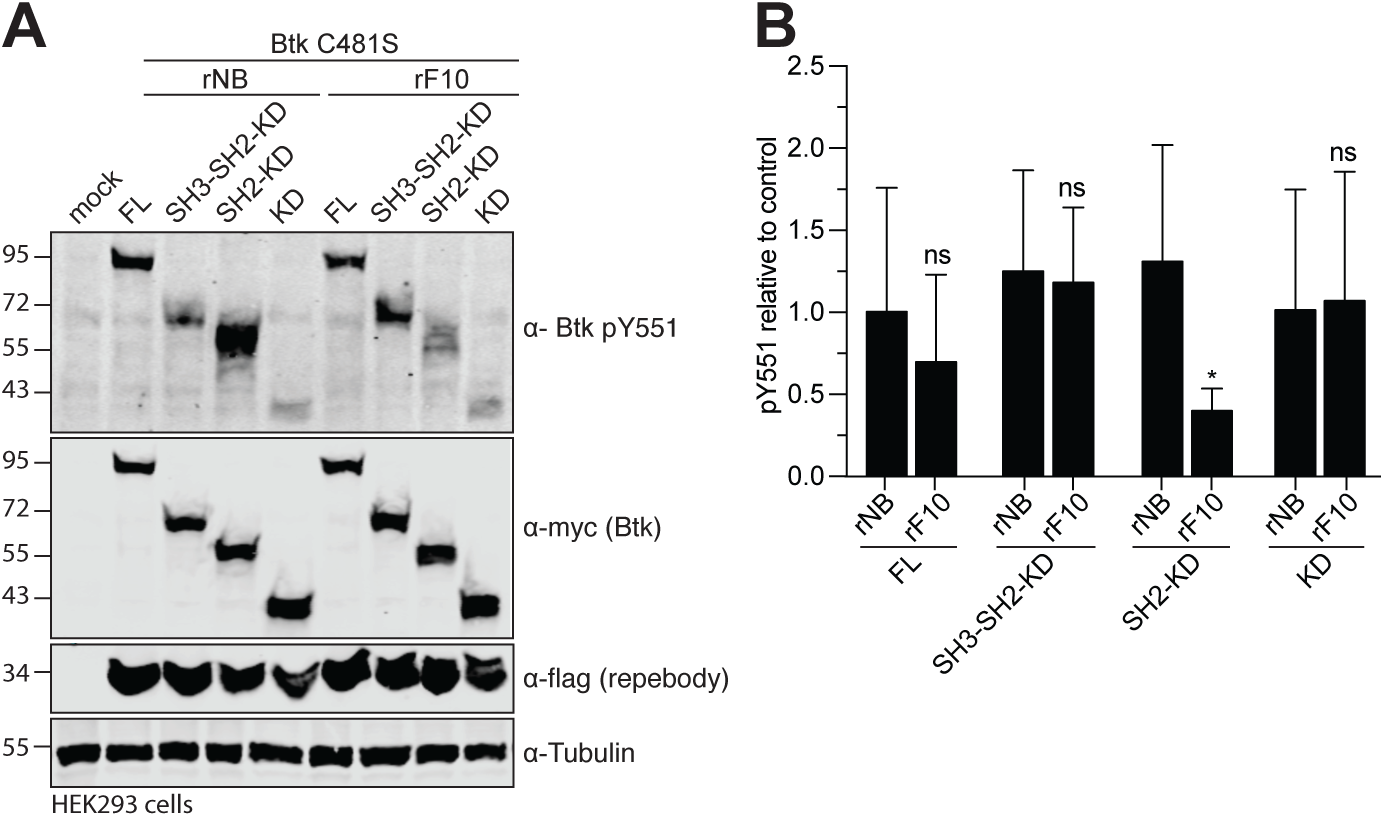
Targeting the Btk SH2-KD interface decreases activation of therapy-resistant Btk with mutation on C481. A. HEK293 cells were transiently transfected with indicated Btk C481S constructs and repebodies. Cell lysates were immunoblotted to assess Btk pY551 phosphorylation. B. Quantification of pY551 shown in (A) and normalized to total Btk expression. Data shown are the mean ± SD of two biological replicates, and P-values were calculated using an unpaired *t*-test. *P ≤ 0.05. See also Figure S7.

To our knowledge, this is the first report of an alternative mechanism able to inhibit wild-type and drug-resistant Btk by targeting an allosteric site. Together, we demonstrated that the SH2-KD interface is critical for Btk kinase activation and a targetable site to improve therapies of Btk-driven malignancies.

## DISCUSSION

While recent studies provided considerable insight on how Btk is autoinhibited, little is known about the interdomain rearrangements coordinating Btk activation (Shah et al., 2018). Here, we demonstrated that an unsuspected allosteric interaction between the Btk SH2 and KD is critical for kinase activation thereby explaining the loss-of-function phenotype of a subset of XLA mutations in the SH2 domain. Furthermore, we identified and validated the SH2-KD interface as a unique site for allosteric Btk inhibition using an engineered repebody protein. To our knowledge, this is the first alternative targeting strategy for ibrutinib-resistant Btk variants. Both FDA-approved Btk inhibitors, ibrutinib and acalabrutinib, are highly susceptible to resistance development by mutations of C481 to which these inhibitors covalently bind. Therefore, the therapeutic exploitation of the Btk SH2-KD interface for patients with TKI-resistant B-cell malignancies is highly attractive. But SH2 domains are conserved, abundant domains in proteomes, and hence difficult to target using small molecules. Small engineered protein binders, in particular monobodies, have emerged as powerful tools to target SH2 domains of a variety of kinases and phosphatases and encourage the development of alternative SH2 inhibitors (Sha et al., 2013, Wojcik et al., 2016, Kukenshoner et al., 2017). In particular, recent advances in cytoplasmic delivery strategies of protein binders, its combination with targeted protein degradation and feasibility PROTAC-based Btk degradation demonstrate progress towards therapeutic applicability (Schmit et al., 2019; Dobrovolsky et al., 2019). The combined targeting of different sites on the same Btk molecule to limit resistance development is likely to follow the paradigm of the allosteric Bcr-Abl myristoyl pocket inhibitor asciminib, which abrogates drug resistance when combined with ATP-competitive Bcr-Abl inhibitors (Eide et al., 2019), In addition to the SH2-kinase interaction, inhibition of dimerization of the PH-TH domain, which is required for membrane-associated activation of Btk, was proposed for allosteric targeting although not yet explored (Wang et al., 2015).

It is important to note that targeting the Btk SH2-KD interface leads to particularly strong effects in DLBCL cell lines harboring CD79B ITAM mutations (Y196F in HBL-1 and Y196H in TMD8 cells) that trigger chronic activation of BCR, and are therefore heavily dependent on Btk signaling to support cell growth. This may indicate that patients with this genotype might benefit strongly from allosteric targeting approaches of Btk (Davis et al., 2010).

Besides B-cell malignancies, Btk is an emerging target in autoimmune diseases, e.g. rheumatoid arthritis. Here, allosteric inhibition of Btk in Fc receptor (FcRs) signaling in basophils and mast cells could provide potential therapeutic benefits (Pal Singh et al., 2018).

Previous studies on other Tec kinase members, in particular Itk, reported an increase in kinase activity in the presence of protein-interacting domains, including the SH2 domain (Joseph et al., 2007, Joseph et al., 2009). This early mechanistically unexplained observation may hint towards conservation of the allosteric SH2-kinase interface in all Tec kinases. Strikingly, residues involved in the Btk SH2-kinase interface are identical in most other Tec members, but are not conserved in the Abl and Fes kinase families and residues critical for the Abl SH2-kinase interaction are not conserved in Tec kinases (Figure S7C; Filippakopoulos et al., 2008, Grebien et al., 2011). This further supports the notion that allosteric SH2-kinase interfaces in different kinases families appear to be diverse in terms of location, charge, hydrophobicity, size, and, importantly, dynamics. The Btk SH2-KD interaction seems highly dynamic, which precluded crystallization, in contrast to the respective Fes and Abl SH2-KD constructs (Filippakopoulos et al., 2008, Lorenz et al., 2015), as well as Csk, where both SH2 and SH3 domains contact the N-lobe (Ogawa et al., 2002).

Our data also provides a first structural insight on the molecular mechanism-of-action of several SH2 mutations in XLA. Those may disturb Btk activity by shifting the SH2-KD conformation towards a more compact and thus less active kinase compared to a native elongated conformation. Additional multi-domain structures including the SH2-kinase linker will potentially capture the preferential position of the SH2 towards the KD, and further corroborate the model of SH2-mediated activation of Tec kinase members.

Our study adds novel structural insights into the complex regulation of Tec kinases, where the SH2 domain plays a critical role in the kinase activation which is independent of its canonical function. Disruption of the SH2-KD interface hampers Btk activation and provide a molecular mechanism that explains a subset of pathogenic XLA mutations. Finally, the exploration of the SH2-KD interface as a targetable allosteric site, even in therapy-resistant Btk variants, provides a completely novel way to target Btk and potentially other Tec kinases.

## SUPPLEMENTAL INFORMATION

Supplemental Information includes seven figures and five tables can be found with the submitted material.

## SUPPLEMENTAL VIDEO

Video 1. Superimposition of active Btk SH2-KD complex with the SH2-rF10 repebody crystal structure (6HTF). See Figure 4H for details.

## ACKNOWLEDGEMENTS

This work was supported by grants from the Swiss Cancer League (grant KLS-3595-02-2015) and the Global Research Laboratory (GRL) Program (Korea National Research Foundation). We thank M. Tully (ESRF Grenoble) for support during SAXS measurements, M. Thome-Miazza for cell lines, A. Reynaud for help with crystallization, L. Menin and D. Ortiz for MS analysis and J. Kuriyan for some Btk expression constructs. We acknowledge the Paul Scherrer Institute (Villigen) for provision of synchrotron radiation beamtime and local contacts for their support during crystal diffraction. We thank all members of the Hantschel lab for continuous support and discussions.

## AUTHOR CONTRIBUTIONS

D.D. conducted and analyzed most experiments. A.L. contributed to study design and experiments. G.L.S. performed the molecular dynamics simulations under M.D.P. and M.D.V. supervision. S.J., Y.-K. S. and H.-S.K. developed the repebody. M.J.M. contributed to the SAXS data acquisition. A.P and D.S. provided assistance for the SAXS analysis. S.G. provided technical assistance and vital tools for all experiments. F.P. performed crystallography data acquisition and support to solve the structure. T.K. assisted with the ITC measurements. D.D. and O.H. designed and coordinated the study, planned the experiments, interpreted the data and wrote the manuscript.

## METHODS

## KEY RESOURCES TABLE

Attached to the Supplemental Information file.

## LEAD CONTACT AND MATERIALS AVAILABILITY

Further information and requests for resources should be directed to and will be fulfilled by the Lead Contact, Oliver Hantschel. Plasmids and cell lines are available upon request to the Lead Contact.

## EXPERIMENTAL MODEL AND SUBJECT DETAILS

### Cell Lines and Culture Conditions

HEK293 and HEK293T cells were cultured in DMEM (Gibco) supplemented with 10% fetal calf serum (Gibco) and 1% penicillin/streptomycin (Bioconcept). Human DLBCL cell lines DOHH2, HBL-1 and TMD8 (expressing full-length wild-type Btk protein) were kindly provided by M. Thome-Miazza (University of Lausanne), cultured in RPMI-1640 media (Gibco) supplemented with 1 mM L-glutamine (Gibco), 10% fetal calf serum (Gibco) and 1% penicillin/streptomycin (Bioconcept). All cell lines were cultured at 37°C under 5% CO_2_.

## METHOD DETAILS

### Protein expression and purification

Btk SH2 domains were cloned into the pETM30 plasmid with an N-terminal 6xHis-GST tag with tobacco etch virus (TEV) cleavage site, and expressed in *E.coli* Bl21(DE3). Repebodies were cloned into the pET21a plasmid (Millipore) with a C-terminal 6xHis tag, and expressed in *E.coli* Origami (DE3). Expression of recombinant proteins was performed overnight at 18°C in LB medium after induction with 0.5 mM IPTG at an optical density of ∼0.8. For protein purification, bacteria were harvested in purification buffer (50 mM Tris pH 7.5, 500 mM NaCl, 1 mM DTT, 5% glycerol, 10 mM imidazole) containing DNAse, homogenized using an Avestin Emulsiflex C3 homogenizer, followed by lysate clarification through centrigugation. Proteins were first purified by gravity flow Ni-NTA agarose (Qiagen, 30210) followed by tag cleavage with recombinant TEV protease in dialysis in buffer (25 mM Tris pH 7.5, 300 mM NaCl, 1 mM DTT, 5% glycerol). Finally, samples were subjected to size exclusion chromatography (SEC) on a Superdex 75 16/60 column equilibrated with dialysis buffer, and peak fractions pooled and analyzed by SDS-PAGE.

For insect cell expression, sequences were cloned into a pFast-Bac-Dual plasmid (Thermofisher). To obtain unphosphorylated Btk, Flag-tagged Yersinia protein tyrosine phosphatase (YopH) was simultaneously expressed from the same vector. Baculoviruses were prepared following the instructions from Bac- to-Bac Baculovirus Expression System (Thermofisher) protocol. Briefly, pFast-Bac Dual plasmids were transfected to *E.coli* DH10B followed by bacmids purification (PureLink, Invitrogen) and transfection in Sf9 cells using the transfection reagent FuGene HD (Promega, E2311). Supernatant containing the baculoviruses were used to produce recombinant proteins in Sf9 cells at density 1.5×10^6^ cells mL^-1^ in SF-900 SFM (10902-096, Thermo) cultured at 28°C and 80% air humidity. After 3 days, cells were and resuspended in purification buffer containing 1mM PMSF, protease cocktail inhibitor (Roche) and Benzonase (Millipore), and lysed by sonication. Cleared lysates were purified and tags removed as described above. All purified proteins could be stored at -80°C without loss of activity. Absence of YopH phosphatase activity was reassured by a phosphatase activity assay using colorimetric PNPP substrate (Thermo) and immunoblotting.

### Site-directed mutagenesis

All point mutations were introduced using the Quikchange II Site-Directed Mutagenesis Kit (Agilent) using primers described in Table S5. Sequence alignments were generated using Geneious (Biomatters).

### Kinase autophosphorylation assay

1 µM of recombinant Btk proteins were incubated in Tris 25 mM pH 7.5, 150 mM NaCl, 5% glycerol, 1 mM ATP, 20 mM MgCl_2_, 1 mM DTT. For inhibition of autophosphorylation, 2 µM of repebodies were pre-incubated with 1 µM of Btk proteins for 15 minutes before starting the reaction upon the addition of 1mM ATP. Reactions were carried out at room temperature and stopped at desired time points by adding 2X Laemmli buffer to each tube, followed by boiling 5 minutes at 95°C. Samples were immunoblotted onto a nitrocellulose membrane using a Dot-Blot apparatus (Bio-Rad).

### HEK293 transfection

Btk constructs were expressed in HEK293 cells using pCS2-gateway plasmid containing an N-terminal 6xMyc tag, while repebodies were cloned into pcDNA3.1 vector and contained a C-terminal Flag tag. Transient transfections with respective plasmids were performed using Polyfect transfection reagent (Qiagen). 48 hours after transfection, cells were harvested, lysed and samples further processed for immunoblotting.

### Cell lysis and immunoblotting

Cells were lysed in IP buffer (50 mM Tris-HCl pH 7.5, 150 mM NaCl, 1% NP-40, 5mM EDTA, 5mM EGTA, 25 mM NaF, 1mM orthovanadate, 1mM PMSF, 10 mg mL^-1^ TPCK and protease cocktail inhibitor from Roche), and cleared by centrifugation at 14,000 rpm for 10 minutes at 4°C. Total protein concentration was measured using Bradford assay (Bio-Rad). All immunoblotting analysis was performed using 100 μg of total protein.

### Antibodies

A list of primary and secondary antibodies is available in the STAR methods table from supplemental material. Western blot quantification of fluorescent secondary antibodies was done using the Li-Cor Odyssey system. ECL prime detection reagent (RPN2232, GE Healthcare) was used to detect HRP-conjugated antibodies using the C-digit Blot scanner (Li-Cor). Western blot normalization was done using total protein signal or loading control (tubulin).

### Immunoprecipitation

Immunoprecipitation from HEK293 cell lysates was done using 1 mg of total protein adjusted to 1 mL volume with IP buffer. Anti-c-Myc Agarose Affinity Gel (Thermo, A7470) was added to the lysates and incubated for 3 hours on a rotating wheel at 4°C. Beads were subsequently washed three times with IP buffer and finally boiled in Laemmli buffer for 5 minutes at 95°C before subjected to immunoblotting.

### Mass spectrometry

For confirmation of protein identity and phosphorylation status, recombinant proteins were analyzed on a Xevo G2-S QTOF mass spectrometer (Waters) operated in positive ionization using the ZSpray™ dual-orthogonal multimode ESI/APCI/ESCi source. Data were processed using MassLynx™ 4.1 software and MaxEnt1 application for deconvolution.

### B cell transduction and BCR stimulation

Repebodies containing a C-terminal flag tag were cloned into the doxycycline-inducible (Tet-ON) lentiviral vector pcW57.1 (Addgene) and co-transfected with envelope and packaging plasmids (pMD2G and pCMVR8.74 respectively, a kind gift from the Trono Lab, EPFL) into HEK293T cells using the CalPhos Mammalian Transfection kit (Clontech). Lentiviruses were concentrated by ultracentrifugation at 30,000xg for 2 hours at 16°C, and added to lymphoma cell lines, followed by a single spinoculation step at 300xg for 60 minutes at room temperature. On the next day, cell media was replaced and cells selected using 2 ug mL^-1^ puromycin. Repebodies were induced by the addition of 2 µg mL^-1^ of doxycycline. For BCR stimulation, cells at density 5×10^6^ cells mL^-1^ were incubated at 37°C with 20 µg mL^-1^ anti-human IgM/G F(ab’)2 from goat (Jackson ImmunoResearch, 109-006-129 and 109-006-098) in RPMI-1640 media without calf serum. Cells were then harvested, immediately lysed and samples further processed for immunoblotting.

### Cell viability assays

DLBCL cell lines were treated for 48 hours with ibrutinib (concentration range 50 nM to 100 µM) and viability assessed using Cell Titer Glow (Promega). Luminescence was measured in a SpectraMax M5 plate reader (Molecular Devices). DMSO (50 µM) and doxorubicin (10 µM) were used as negative and positive controls, respectively. To assess the effects of repebodies on the viability of DLBCL, transduced cells were treated with 2 µg mL^-1^ of doxycycline to induce expression of repebodies, and cell number verified using a Casy Cell Counter (OLS Omni Life Science). Cell density was maintained at 5×10^5^ cells mL^-1^ and regularly diluted when cell density reached 3×10^6^ cells mL^-1^. Parental (non-transduced) and non-induced cells were used as control.

### Molecular dynamics (MD) simulations

System preparation: The Btk SH2-KD structural model used for our MD simulations was built starting from the X-ray structures of the apo-KD of Btk (PDB: 1K2P) and SH2 domain (PDB: 6HTF). The SH2 domain was initially positioned on the top of KD at ∼60 Å from it (the distance is measured considering the center of mass of SH2 and N-lobe of KD). The linker sequence was manually added by means of Maestro (Schrödinger Release 2016-1: Maestro, Schrödinger, LLC, New York, 2016) obtaining an extended configuration (Fig. 3A) and subsequently refined through a scaled MD simulation run. Firstly, the protein was parameterized by using the Amber 14SB force field (Maier et al., 2015) and immersed in a TIP3P (Jorgensen and Madura, 1983) water box having 12 Å of buffer between the protein and the three edges of the box. The protein was then neutralized by adding an appropriate number of Cl^-^ ions. After minimization, the protein underwent to three NVT simulations steps of 500 ps each to gradually reach the target temperature of 300 K (first step from 0 to 100 K, second step from 100 K to 200 K and last step from 200 K to 300 K). Here, a restraints of 1000 kJ mol^−1^ nm^−2^ was applied to backbone and the velocity-rescaling thermostat (Bussi et al., 2007) was used. Then, 1 ns of NPT simulation was performed maintaining the restraints and employing the Parrinello-Rahman barostat (Parrinello and Rahman, 1981) to reach the target pressure of 1 bar. Finally, to enhance the sampling of the linker sequence, we performed a 70 ns long Scaled MD simulations (Mollica et al., 2015b), using a λ = 0.8, and releasing the restraints for the linker sequence. Electrostatics were treated with the cutoff method for short-range interactions and with the Particle Mesh Ewald method (Darden et al., 1993) for the long-range ones (rlist= 1.1 nm, cutoff distance= 1.1 nm, VdW distance = 1.1 nm, PME order= 4). To obtain the optimal configuration of the linker sequence, we performed a cluster analysis on the last 70 ns long trajectory. The centroid of the most populated cluster was then employed as starting point for the next MD simulations. Both the MD simulations and cluster analysis were performed by using BiKi LifeSciences suite. (Decherchi et al., 2018).

Scaled MD simulations: We run multiple replicas Scaled MD simulations (Mollica et al., 2015b) to enhance the sampling of Btk kinase and speed up the binding events between SH2 and KD. As starting point, we employed the refined structure of the Btk SH2-KD-linker model (see system preparation section). The equilibrated system was submitted to 40 replicates ∼100 ns long Scaled MD simulations using a λ = 0.9. Here, restraints were not applied because the high scaling factor enabled adequate sampling without affecting the overall folding of the system. BiKi LifeSciences suite (Decherchi et al., 2018) was used for Scaled MD simulations, using the same settings as in the system preparation section.

Data Analysis: The final aim of our MD studies is to collect all possible Btk SH2-KD bound configurations and to determine which is the most likely bound configuration(s) using both SAXS and mutational studies data. The collected 4 µs-long scaled MD trajectories resulted in a total of 400,000 frames. From this large ensemble, we extracted the unique and non-redundant Btk SH2-KD configurations, using a clustering procedure implemented in BiKi (Decherchi et al., 2018). The resulting 760 structures were submitted to another cluster analysis to probe the preferred 3D structural organization of SH2 domain with respect of KD (Fig. 3A and S3A). Also, we run a CRYSOL analysis (Svergun et al., 1995) to compute the χ^2^ value for each structure, in order to select the SH2-KD complexes with the best fitting with the experimental SAXS curves.

### Development of repebodies

Selection and affinity maturation of human Btk SH2-specific repebody (rF10) were performed through phage display and a modular evolution approach as previously described (Lee et al., 2012).

### Small-angle X-ray scattering (SAXS)

SAXS data were collected at the BM29 beamline (ESRF Grenoble, France). All proteins were measured in buffer containing 25mM Tris pH 7.5, 300 mM NaCl, 1 mM TCEP and 5% glycerol. A robotic sample changer carried out the measurement in the batch mode, while in-line SEC-SAXS was performed using a Superdex S200 Increase 10/300 column (GE Healthcare) with a flow rate of 0.7 mL min^-1^ at room temperature. Acquired data were averaged and subtracted from an appropriate solvent-blank to produce the final curve using the ATSAS Suite, EMBL (Franke et al., 2017) and CHROMIXS (Panjkovich and Svergun, 2016a). Initial data pre-processing and reduction were performed using an automatic pipeline. Final scattering curves were analyzed using PRIMUS for evaluation of molecular dimensions *(R_g_)* (Konarev et al., 2003) and maximum particle dimension (D_max_) using GNOM (Svergun, 1992). Moreover, the Porod volume was computed using the Porod invariant (Porod, 1952), and the molecular mass estimated using SAXSMoW 2.0 (Fischer et al., 2010), Bayesian inference approach (Hajizadeh et al., 2018) and Volume-of-correlation (Rambo and Tainer, 2013). Data collection and summary of structural parameters are described in Table S2. *Ab initio* models were computed with DAMMIF (Franke and Svergun, 2009). SREFLEX (Panjkovich and Svergun, 2016a) was employed to improve the agreement of flexible multidomain models to the experimental data. Finally, the flexibility of multidomain complexes was assessed with Ensemble Optimization Method 2.0 (Tria et al., 2013). Fitting of models to experimental data was assessed using CRYSOL (Svergun et al., 1995) molecular and superimpositions performed with the SASpy (Panjkovich and Svergun, 2016b). Data collection and structure determination statistics are described in Table S2 and S4.

### Crystallization, data collection, and structure determination

Recombinant Btk SH2 and rF10 proteins were mixed at 1:1 ratio and the complex purified with a Superdex 75 column 16/600 (GE Healthcare) in buffer containing 25 mM Tris pH 7.5, 300 mM NaCl, 1 mM TCEP. The purified complex was concentrated to ∼25 mg mL^-1^ and crystallized at 18°C using the hanging-drop vapor-diffusion method by mixing 1:1 with a solution containing 1 M Tris pH 8.5, 300 mM sodium fluoride, 300 mM sodium bromide, 300 mM sodium iodide, 25% MPD; 25% PEG 1000; 25% PEG 3350. 20% glycerol was used as a cryoprotectant. X-ray diffraction data was collected at the SLS Beamline X06DA in the Swiss Lightsource (SLS, Villigen, Switzerland) at a wavelength of 1 Å and temperature of 100 K. Data collection and structure determination statistics are described in Table S3. Diffraction data was processed and scaled with the XDS package. The structure was solved by molecular replacement employing models derived from a previously reported repebody (PDB 5B4P) and Btk SH2 (PDB 2GE9) excluding loop regions. Molecular replacement, manual model building, B-factor refinement, solvent addition, energy-minimization and refinement of structures were conducted iteratively using Phaser and Coot (Phenix version 1.13). Molecular graphics were generated using PyMOL (DeLano Scientific).

### Isothermal titration calorimetry (ITC)

Proteins were extensively dialyzed in buffer containing 20 mM Hepes pH 7.5 and 150 mM NaCl, briefly degassed, and concentration determined by measuring UV absorbance at 280 nm. ITC measurements were performed on a MicroCal PEAQ-ITC (Malvern) instrument. The repebody (100 µM) was titrated into SH2 domains (10 µM) at room temperature in 16 steps with 0.49 μL for the first and 2.49 μL for the other steps. Thermodynamic parameters were obtained using the MicroCal software.

### Fluorescent Polarization (FP) binding assays

Btk SH2 WT and mutants were incubated at several concentrations with 1µM of FITC-labeled ADNDpYIIPLPD peptide in Tris 40 mM pH 8, 150 mM NaCl and 1 mM DTT. Competitive FP assay was performed using 25 µM of Btk SH2 and 1 µM of peptide incubated with repebody in a range of 200 µM - 20 nM. FP signal was measured using a SpectraMax M5 plate reader (Molecular Devices) with excitation at 485 nm and emission at 530 nm in a 96 well black-plate (Greiner, 784-900).

### Multi-angle light scattering analysis (SEC-MALS)

Multi-angle light scattering was used to probe for oligomerization states. All measurements were performed at room temperature using a Dawan Hellios multi-angle light scattering detector (Wyatt Technologies) coupled to an SEC column. 80 μL (0.5 mg mL^-1^) of purified recombinant protein was injected into a Superdex 75 HR10/30 column (GE Healthcare) in buffer containing 25 mM Tris-HCl pH 7.5, 150 mM NaCl, 1 mM TCEP, and eluted at a flow rate of 0.5 mL min^-1^. Absolute molecular weight and homogeneity were determined using ASTRA version 5.3 (Wyatt Technologies).

### Circular dichroism (CD)

Far-UV spectra (190-300 nm) of recombinant Btk SH2 WT and mutants were carried out in buffer containing 10 mM Na-phosphate buffer pH 7.2 and 100 mM NaF using a 0.1 cm quartz cell and CD Spectrometer Chirascan V100 (AppliedPhotophysics). Data was acquired at a step size of 1 nm and bandwith of 1 nm. 3 scan records for each protein were subtracted from the background (buffer only) and averaged to generate the data reported in units of mean molar ellipticity per residue. Melting curve analysis was performed by measuring proteins at the wavelength corresponding to the peak for the predominantly ß-sheet SH2 domain (218 nm) in a temperature range from 20 to 94°C, ramp-rate 1°C per minute.

### *In vitro* Kinase assay

2 µM of repebodies were pre-incubated with 1µM of recombinant Btk proteins in buffer Tris 25 mM pH 7.5, 150 mM NaCl, 5% glycerol, 20 mM MgCl_2_, 1 mM DTT in the presence of 50 μM ATP, 7 μCi γ-^32^P-ATP, and PLCγ2 peptide carrying an N-terminal biotin (biotin-ERDINSLYDVSR-amide). Peptide concentrations ranged from 100 μM to 3.125 μM. Reactions were carried out on a final volume of 20µL at room temperature for 20 minutes and terminated using 10 μL 7.5 M guanidinhydrochlorid. Samples were spotted onto a SAM2 Biotin Capture membrane (Promega) and further treated according to the instructions of the manufacturer.

### Mapping of Btk autophosphorylation sites

Sample preparation: Recombinant autophosphorylated Btk SH2-KD was separated by SDS-PAGE and stained with Coomassie. Bands of interest were excised, in-gel digested in reduced in 10 mM DTE, 50 mM AB, and then alkylated in 55 mM iodoacetamide, 50 mM AB. After a washing step, gel extracts were digested with MS Grade Trypsin over-night. Resulting peptides were finally extracted using a high organic containing solvent and dried by vacuum centrifugation prior to LC-MS2 measurements or phosphopeptides enrichment.

Next, 90% of the extracted peptide was used for phosphopeptides enrichment step while the remaining 10% was used for sample identification. Titanium dioxide affinity principle was used for enrichment using home-made titania tips (based on Thingholm and Larsen 2009). Dried samples were resuspended 0.75% TFA, 60% acetonitrile, 300 mg ml^-1^ lactic acid, loaded on tips, and eluted in 0.5% ammonium hydroxide and 5% piperidine. Samples were acidified and dried down prior to LC-MS2 measurements.

MS analysis: For the MS detection of phosphopeptides, dried samples were resuspended in 0.1% TFA and separated by C18 Reverse Phase nano UPLC using a Dionex Ultimate 3000 RSLC system (Thermo Fischer) connected to an Orbitrap Elite Mass Spectrometer (Thermo Fischer). Samples were first trapped on a home-made capillary C18 pre-column and then separated on a C18 capillary column (Nikkyo Technos Co; Magic AQ C18; 3 µm - 100 Å; 15 cm x 75 µm ID). Data-dependent mode was used for MS acquisitions were the 20 most intense parent ions were selected for subsequent fragmentation by CID. A potential phosphopeptides m/z inclusion list was also generated and used to maximize detection chances.

## QUANTIFICATION AND STATISTICAL ANALYSIS

All data reported were analyzed using Prism 7 (GraphPad) using software-defined fitting models and unpaired *t*-test statistical test. Calculated P-values are indicated as non-significant (ns), p ≤ 0.05 (*), p ≤ 0.01 (**), p ≤ 0.001 (***) and p ≤ 0.0001 (****).

## DATA CODE AND AVAILABILITY

The X-ray structure of the rF10-SH2 complex was deposited at <u>Protein Data Bank</u> (entry 6HTF). Full SAXS curves and analyzed data for wild-type Btk proteins were deposited at SASBDB (entries SASDF53, SASDF63, SASDF73, and SASDF83).

