## Supplemental Material for "Btk SH2-kinase interface is critical for allosteric kinase activation and its targeting inhibits B-cell neoplasms"

**Content**

**Supplemental Figures S1-S7**
**Supplemental Tables S1-S5**
**Key resources Table**
**Supplemental Video 1**

### SUPPLEMENTAL FIGURES S1-S7

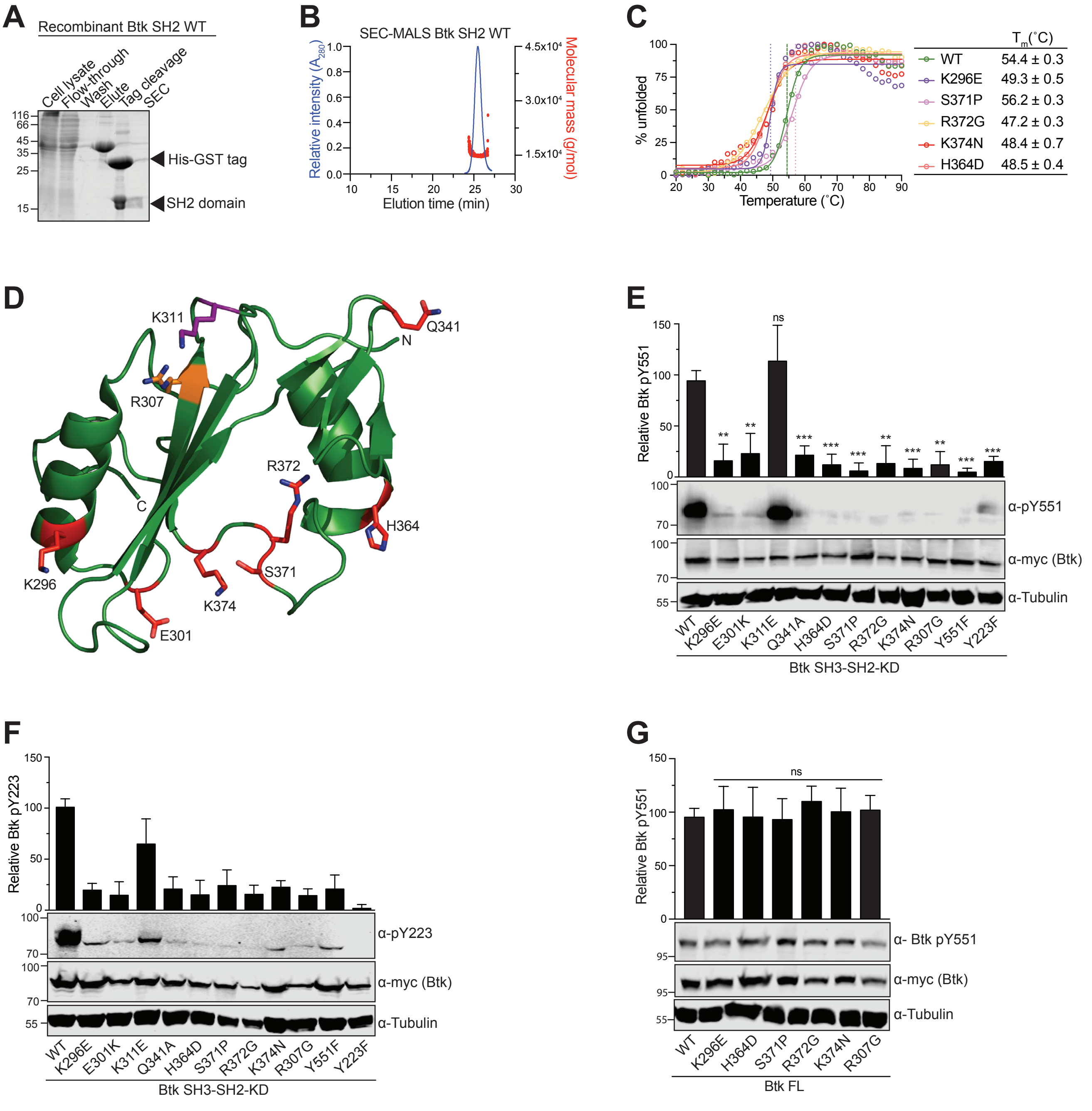


**Figure S1, related to Figure 1. Effect of XLA mutations *in vitro* and in HEK cells.**
(A) Representative SDS-PAGE analysis of purification steps for recombinant Btk SH2 wild-type from *E.coli*. TEV cleavage was used to removal of 6xHis-GST tag used for purification. All Btk SH2 mutants were purified with a similar protocol.
(B) Representative SEC-MALS analysis of purified Btk SH2 domain (monomer = 13.2 kDa). All proteins were analyzed by SEC-MALS and found in the homogenous state in solution (data not shown).
(C)Thermal shift assay (TSA) of recombinant Btk SH2 domains. Melting temperature (Tm) for wild-type Btk was calculated from two independent measurements.
(D) Mapping of a subset of XLA-patient mutations (red sticks) onto the human Btk SH2 structure (PDB 2GE9). The residue R307 (orange sticks) is part of the pY-binding motif (FIVRD). The residue K311 is a non-XLA control mutation facing the opposite surface of the SH2 domain. N- and C-terminal are indicated as N and C, respectively.
(E/F/G) HEK293 cells were transiently transfected with indicated Btk constructs containing an N-terminal 6xmyc tag. Immunoblotting of total cell lysates was performed to assess Btk phosphorylation on sites Y551 and pY223, and relative phosphorylation normalized to total Btk (Myc-Btk) expression. Tubulin was used as loading control. Data shown in E and G are the mean ± SD of three biological replicates, while data shown in F is the mean ± SD of two technical replicates. P-values were calculated against the wild-type (WT) using an unpaired *t*-test. **P ≤ 0.01, ***P ≤ 0.001, and non-significant (ns).

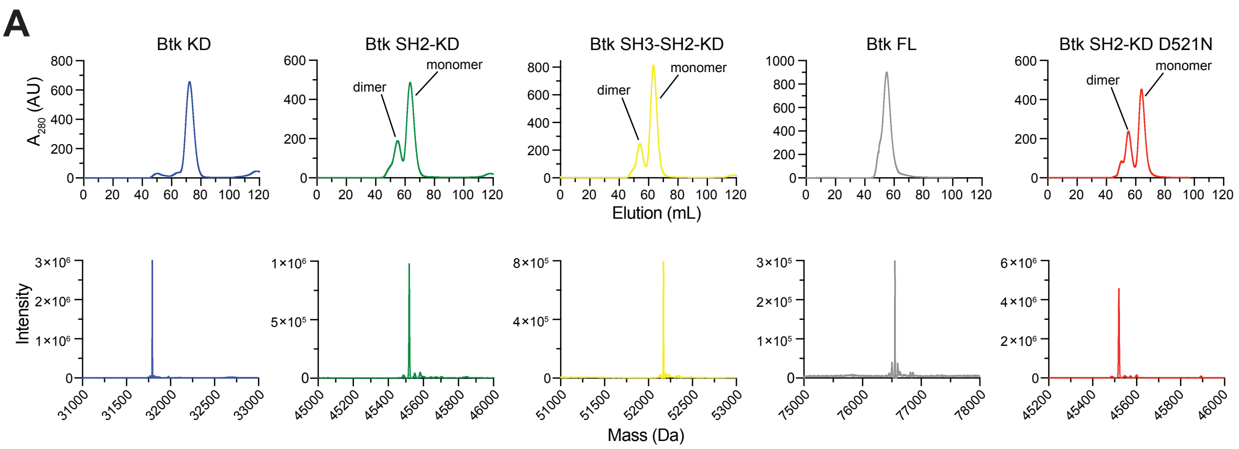


Mass spectrometry analysis

| | Btk KD | Btk SH2-KD (monomer) | Btk SH3-SH2-KD (monomer) | Btk FL | Btk SH2-KD D521N (monomer) |
|---|---|---|---|---|---|
| Obtained (Da) | 31794.09 | 45520.35 | 52171.90 | 76553.45 | 45520.19 |
| Theoretical (Da) | 31793.52 | 45519.93 | 52171.42 | 76553.51 | 45518.95 |

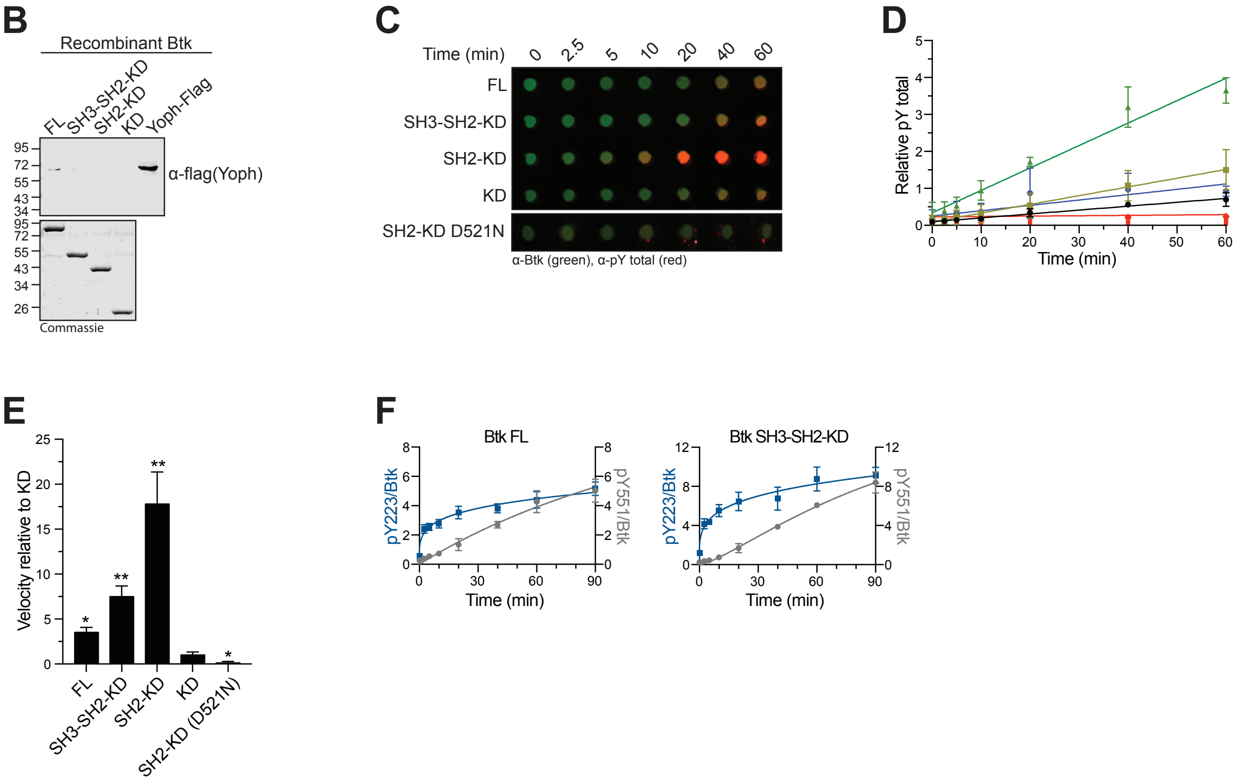


**Figure S2, related to Figure 2. Purification of recombinant Btk and autophosphorylation *in vitro*.**
(A) SEC of recombinant Btk expressed and purified from Sf9 cells (top). Only monomeric peaks were used for the described assays. All samples were subjected to MS analysis for confirmation of protein identity and unphosphorylated state (> 95% for all samples, bottom). The table summarizes the theoretical and obtained molecular weight for the indicated proteins.
(B) Representative immunoblot to confirm the absence of recombinant Yoph-flag phosphatase from proteins purified in Sf9 cells (top) and corresponding SDS-PAGE of recombinant untagged Btk proteins (bottom).
(C/D/E) Btk autophosphorylation *in vitro* assay performed as described in methods. The levels of total phosphotyrosine and total Btk were assessed using immunoblot in a dot-blot apparatus, relative autophosphorylation kinetics plotted overtime and normalized to total Btk protein, and relative autophosphorylation velocities. Data are the mean ± SD of three independent experiments. P-values relative to Btk KD were calculated using unpaired *t*-test. *P ≤ 0.05 and **P ≤ 0.01.
(F)Autophosphorylation kinetics of Y223/Y551 *in vitro* was assessed as described above. Data shown are the mean ± SD of two independent experiments done in duplicates.

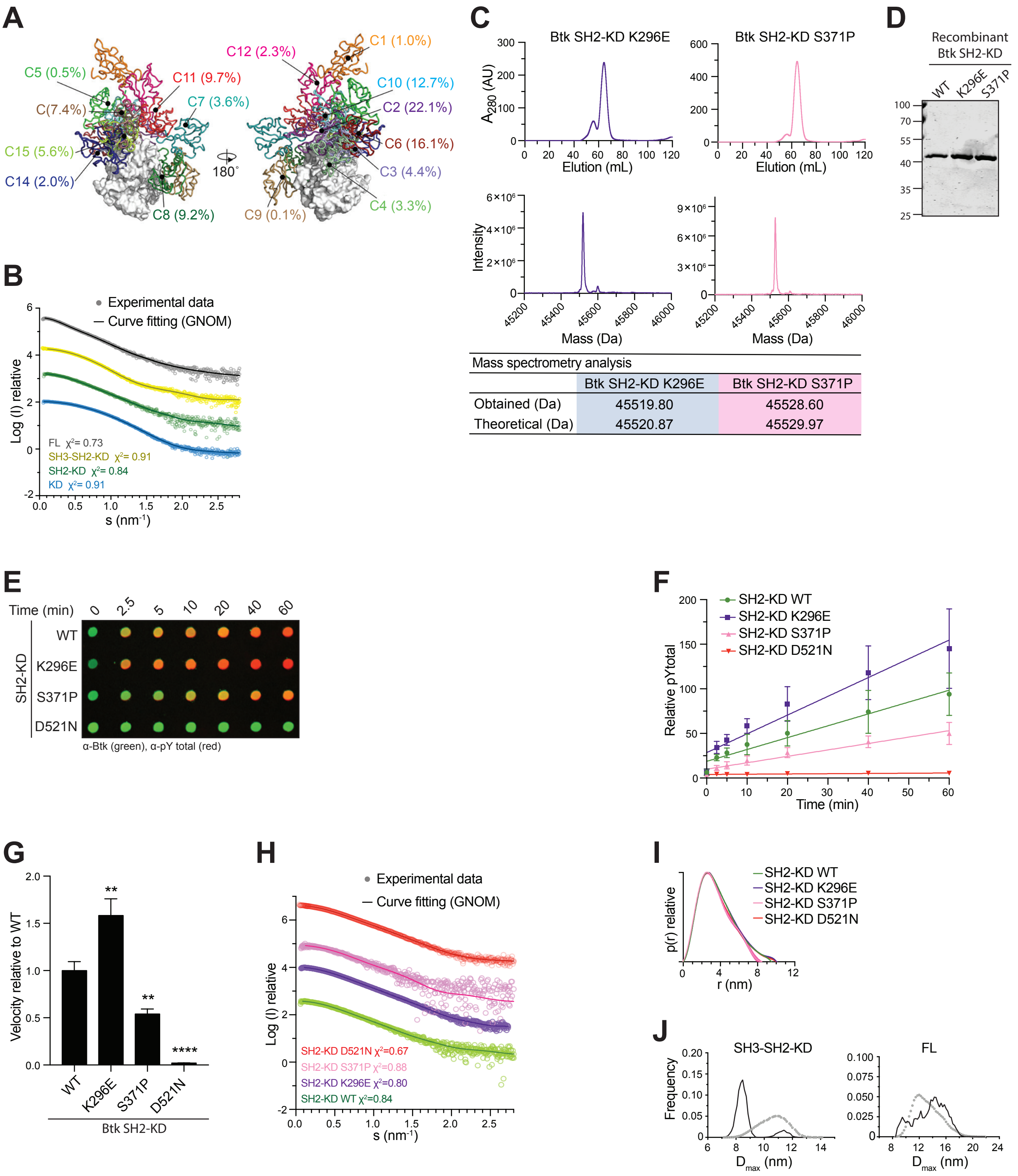


**Figure S3, related to Figure 3. MD and SAXS analysis of Btk wild-type and mutants.**
(A) MD simulation for the SH2-KD complex. Obtained clusters of the SH2 positions (several colors) relative to the KD (white). The percentages indicate the population of the cluster with respect to the entire simulation time (i.e., 4 µs).
(B) Experimental SAXS data of recombinant wild-type Btk proteins. The indicated $\chi 2$ represents the GNOM fitting (line) against the experimental data (dots) for each respective construct. See table S2 for details. Raw curves and full analysis are available in the SASDB database.
(C) SEC of recombinant mutant Btk purified from Sf9 cells (top). All samples were subjected to MS analysis for confirmation of protein identity and unphosphorylated state (>95% for all samples, bottom).
(D) Representative SDS-PAGE analysis of recombinant untagged Btk SH2-KD mutant proteins purified from Sf9 cells.
(E/F/G) Autophosphorylation *in vitro* assay of Btk SH2-KD mutants performed as described in methods. The levels of total phosphotyrosine and total Btk were assessed using immunoblot in a dot-blot apparatus, relative autophosphorylation kinetics plotted overtime and normalized to total Btk protein, and relative autophosphorylation velocities. Data are the mean ± SD of three independent experiments. P-values relative to Btk wild-type (WT) were calculated using unpaired *t*-test. **P ≤ 0.01, ***P ≤ 0.001 and ****P ≤ 0.0001.
(H) Experimental SEC-SAXS data of mutant Btk proteins as indicated in (B).
(I) Maximal particle dimension ($D_{max}$) of mutant Btk proteins. See Table S2 for details.
(J) Flexibility analysis (EOM 2.0) of indicated wild-type Btk proteins showing the $D_{max}$ of selected conformers (lines) from a representative pool of theoretical conformations (dot line).

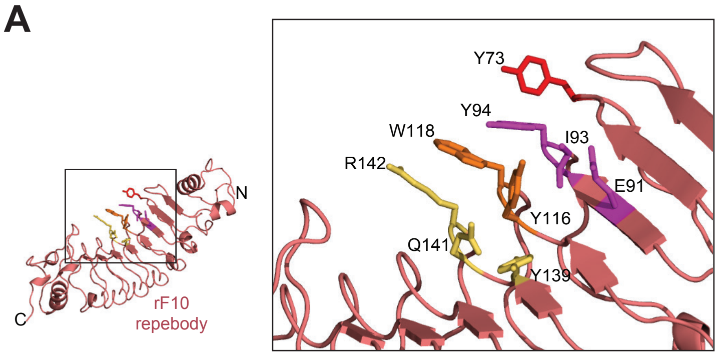


Diversified residues in the Btk SH2 repebody (rF10)

| | LRRV1 | LRRV2 | | | LRRV3 | | LRRV4 | | | $K_d$ (nM) |
|---|---|---|---|---|---|---|---|---|---|---|
| Position | 73 | 91 | 93 | 94 | 116 | 118 | 139 | 141 | 142 | |
| rNB (template) | N | I | T | G | V | E | N | A | H | N.B. |
| rF10 | Y | E | I | Y | Y | W | Y | Q | R | 15.5±1.3 |

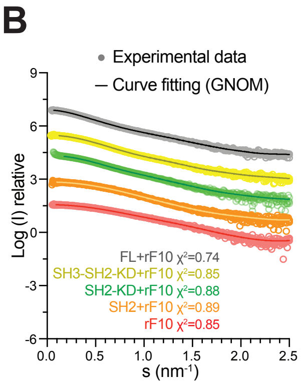


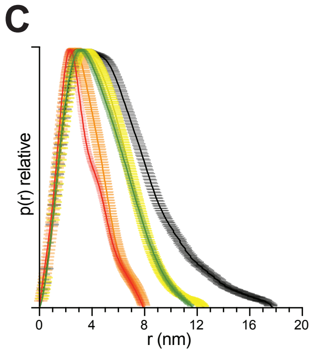


Structural parameters obtained from SAXS

| | rF10 | rF10+SH2 | SH2-KD+rF10 | SH3-SH2-KD+rF10 | FL+rF10 |
|---|---|---|---|---|---|
| $R_g$(nm) | 2.42±0.2 | 2.6±0.14 | 3.48±0.17 | 3.55±0.26 | 4.48±0.38 |
| $D_{max}$(nm) | 7.9±0.8 | 8.2±0.8 | 11.5±1.2 | 12.5±1.2 | 16.9±1.6 |

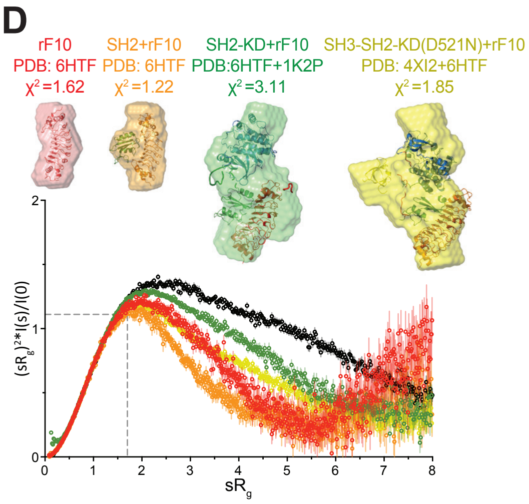


**Figure S4, related to Figure 4. rF10 development and SAXS analysis of rF10-Btk complexes.**
(A) The rF10 repebody (cartoon representation, salmon) was developed by randomizing variable sites within leucine-rich repeats (LRRV) using phage display and modular evolution approach. The residues from the LRRV1 (red), LRRV2 (magenta), LRRV3 (orange) and LRRV4 (yellow) mediating the binding to Btk SH2 domain are indicated as sticks. The table shows the amino acid sequence and binding affinity to the human Btk SH2 domain. Non-binding (N.B.).
(B) Experimental SEC-SAXS data for rF10 alone and rF10-Btk complexes. The indicated χ2 represents the GNOM fitting (line) against the experimental data (dots) for each respective sample. See Table S4 for details.
(C) Maximal particle dimension ($D_{max}$) of rF10 alone and rF10-Btk complexes. The table summarizes the particle dimensions ($R_g$ and $D_{max}$) and the ± error for the indicated constructs.
(D) Dimensionless Kratky plot of rF10 alone and rF10-Btk complexes. *Ab initio* reconstructions obtained from SAXS (surface representation) were superimposed to the indicated crystal/MD structures. For the rF10-SH2-KD and rF10-SH3-SH2-KD complexes, rigid body modeling using SASREF was applied to obtain the final models displayed.

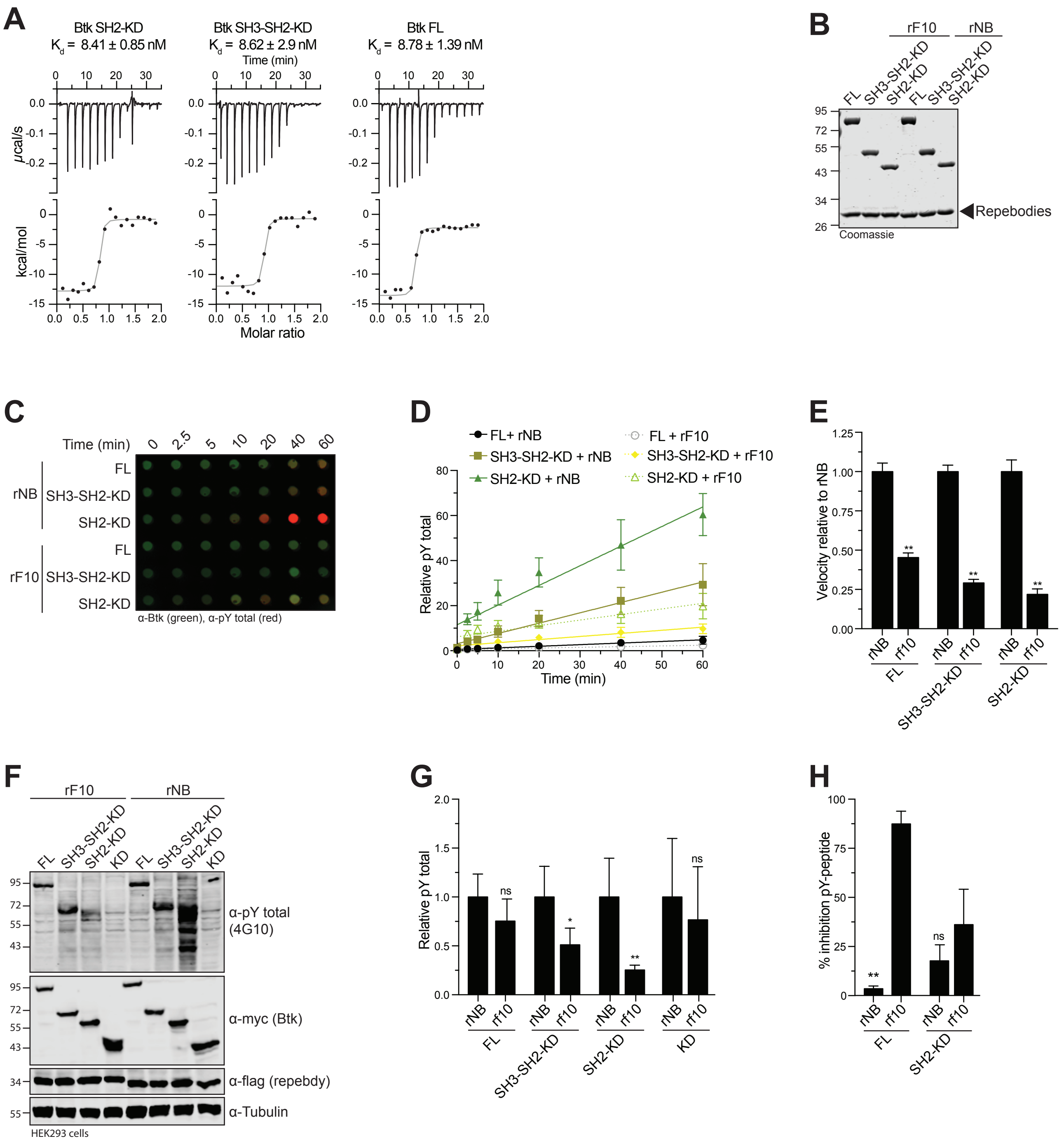


**Figure S5, related to Figure 5. Functional characterization of the rF10 repebody.**

(A) ITC measurement of rF10 repebody to different Btk constructs containing the SH2 domain (SH2-KD, SH3-SH2-KD and FL proteins). Top panels show the raw signal from a representative measurement, and bottom panels show the integrated calorimetric data of the area of each peak. The continuous line indicates the best fit to the experimental data assuming a 1:1 binding model. The $K_d$ (± SD) value was calculated from two independent measurements.

(B) Representative SDS-PAGE analysis of recombinant Btk proteins mixed with rF10 and rNB control repebodies and used for autophosphorylation inhibition *in vitro*.

(C/D/E) Autophosphorylation assay for Btk proteins in the presence of indicated repebodies was performed as described in methods. The levels of total phosphotyrosine and total Btk were assessed using immunoblot in a dot-blot apparatus. Relative autophosphorylation kinetics in the presence of rF10 (dashed lines) or rNB (continuous lines) repebodies plotted overtime and normalized to total Btk protein, and relative autophosphorylation velocities relative to each control repebody. Data are the mean ± SD of three independent experiments. P-values relative to each control rNB repebody were calculated using unpaired *t*-test.

(F/G) HEK293 cells were transiently co-transfected with indicated Btk constructs and repebodies. Immunoblot was used to assess total phosphotyrosine phosphorylation. Quantification of total phosphotyrosine normalized to total Btk (Myc-Btk) expression level and relative to control repebody. Data shown are the mean ± SD of three biological replicates, and P-values were calculated against each control rNB repebody using unpaired *t*-test.

(H) Btk kinase activity against a PLCγ2 peptide (ERDINSL$_{753}$YDVSR) in the presence of rF10 or control repebody. Reported inhibition (% of inhibition of peptide phosphorylation) from two independent experiments done in triplicates. P-values were calculated using unpaired *t*-test. *P ≤ 0.05, **P ≤ 0.01, and non-significant (ns).

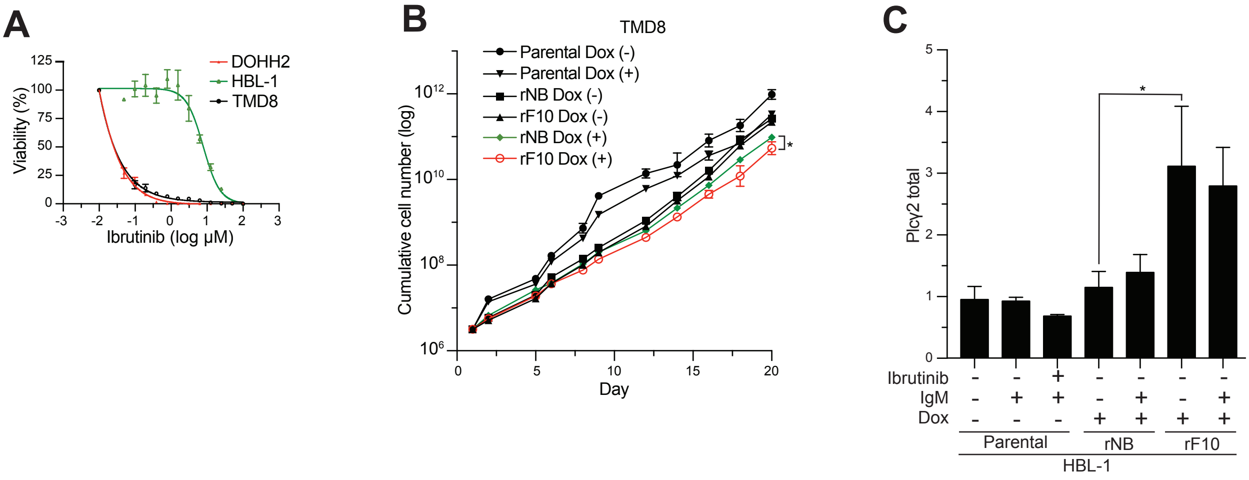


**Figure S6, related to Figure 6. Effect of rF10 in DLBCL cell lines.**

(A) Dose-response for ibrutinib in human DLBCL cell lines. Data points represent the mean ± SD of a representative experiment done in triplicates. Cellular IC50 were obtained by non-linear regression curve fit analysis.

(B) DLBCL TMD8 cell line was transduced with a doxycycline-inducible system for expression of repebodies, and cumulative cell number monitored upon treatment with 2 μg $mL^{-1}$ of doxycycline. Parental cells are non-transduced cells.

(C) Quantification of total PLCγ2 level from HBL-1 inducibly expressing repebodies (flag tagged) for 48 hours. BCR stimulation and ibrutinib treatment were performed as described in methods. Data shown are the mean ± SD from two biological replicates, and P-values were calculated using unpaired *t*-test. *P ≤ 0.05.

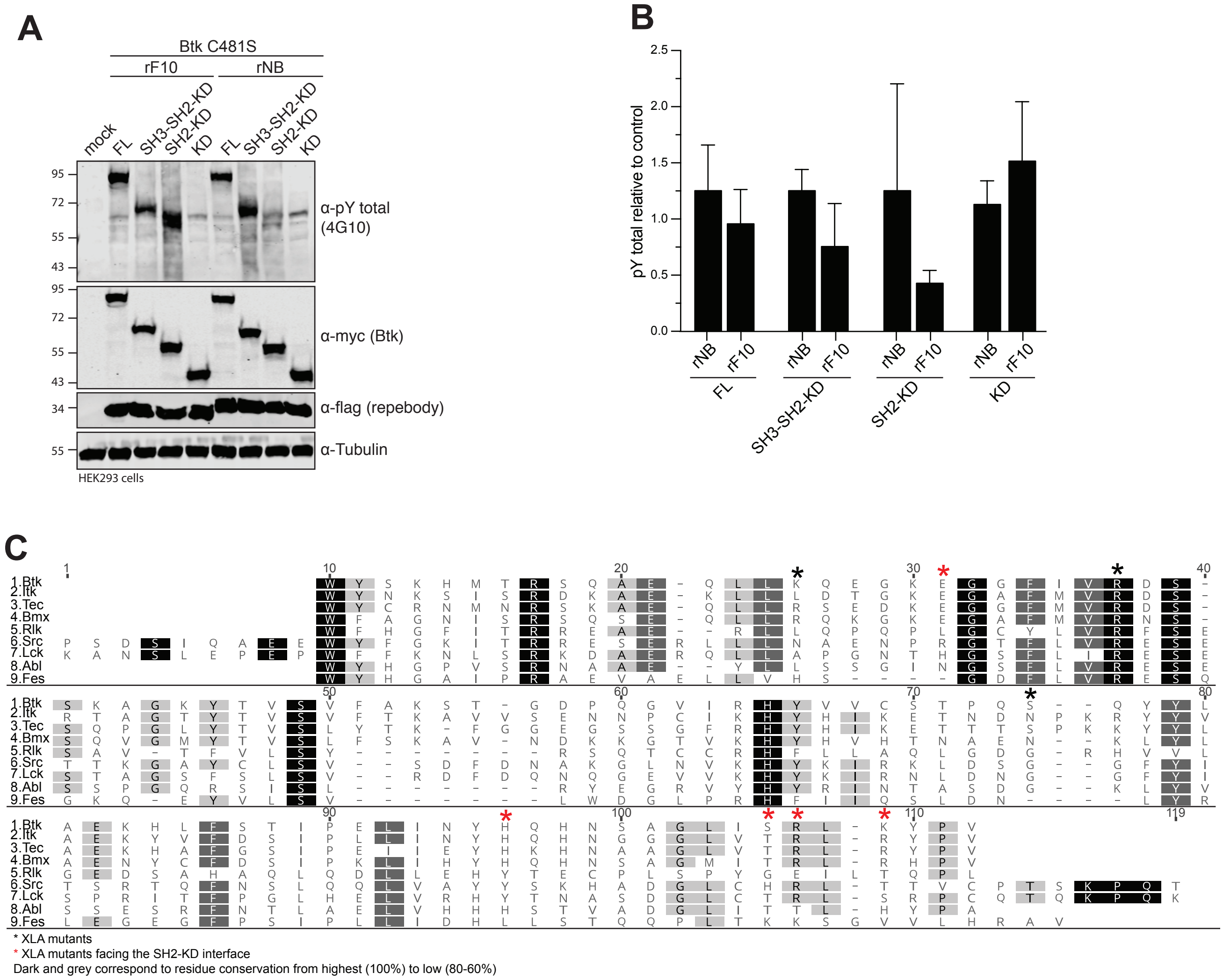


**Figure S7, related to Figure 7 and Discussion. Targeting the Btk SH2-KD interface decreases activation of therapy-resistant Btk with mutation on C481.**

(A/B) HEK293 cells were transiently co-transfected with indicated Btk-C481S constructs and repebodies. Immunoblot was used to assess total phosphotyrosine phosphorylation. Quantification of total phosphotyrosine normalized to total Btk (myc-Btk) expression level and relative to control repebody. Data shown are the mean ± SD of two biological replicates.

(C) Sequence alignment of SH2 domains from human Btk and related kinases. Black squares represent residues highly conserved and grey squares show residues relatively conserved. Black asterisk symbol (*) indicates residues mutated in XLA patients, while red asterisks are residues mutated in XLA patients and located in the predicted SH2-KD interface.

### SUPPLEMENTAL TABLES S1-S5

Table S1, related to Results. *In vitro* autophosphorylation sites on the Btk SH2-KD protein.

| Site | Peptide sequence | Location | Peptide count |
|---|---|---|---|
| Y279 | SDSIEMYEWYSK | SH2 | 56 |
| Y282 | SDSIEMYEWYSK | SH2 | 56 |
| Y315 | AGKYTVSVFAK | SH2 | 60 |
| Y334 | STGDPQGVIRHYVVCSTPQSQYYLAEK | SH2 | 46 |
| Y344 | STGDPQGVIRHYVVCSTPQSQYYLAEK | SH2 | 46 |
| Y345 | STGDPQGVIRHYVVCSTPQSQYYLAEK | SH2 | 46 |
| Y361 | HLFSTIPELINYHQHNSAGLISRLK | SH2 | 4 |
| Y375 | YPVSQQNK | SH2-KD linker | 3 |
| Y392 | NAPSTAGLGYGSWEIDPK | SH2-KD linker | 10 |
| Y425 | WRGQYDVAIK | KD | 10 |
| Y461 | LVQLYGVCTK | KD | 2 |
| Y511 | DVCEAMEYLESK | KD | 4 |
| Y545 | VSDFGLSRYVLDDEYTSSVGSK | KD | 37 |
| Y551 | VSDFGLSRYVLDDEYTSSVGSK | KD | 37 |
| Y571 | FPVRWSPPEVLMYSK | KD | 15 |
| Y627 | VYTIMYSCWHEK | KD | 6 |
| Y631 | VYTIMYSCWHEK | KD | 6 |
| | Total peptide count | | 444 |

Table S2, related to Figure 3. SAXS parameters of human Btk wild-type and mutants.

| **Data collection parameters** | | | | | | | |
|---|---|---|---|---|---|---|---|
| Instrument | MD29 beamline, ESRF Grenoble - France | | | | | | |
| Wavelength (Å) | 0.9919 | | | | | | |
| q-range ($nm^{-1}$) | 0.03563 - 5 | | | | | | |
| Exposure time (sec) | 5 (10 frames x 0.5 sec) | | | | | | |
| Temperature (K) | 290 | | | | | | |
| **Samples** | **KD** | **SH2-KD** | **SH3-SH2-KD** | **Full-length** | **SH2-KD K296E** | **SH2-KD S371P** | **SH2-KD D521N** |
| Measurement mode | batch | SEC-SAXS | batch | batch | SEC-SAXS | SEC-SAXS | batch |
| Concentration range ($mg.ml^{-1}$) | 0.9 – 4.2 | 100μ at 20.4 | 0.9 – 9.2 | 0.4 – 5.5 | 100μl at 30 | 100μl at 10.8 | 0.6 – 1.9 |
| SASBDB identifier | SASDF53 | SASDF63 | SASDF73 | SASDF83 | N/A | N/A | N/A |
| **Structural parameters** | | | | | | | |
| $R_g$ (nm) from Guinier | 2.09 ± 0.02 | 2.832 ± 0.3 | 2.62 ± 0.15 | 4.04 ± 0.03 | 2.92 ± -0.04 | 2.64 ± 0.23 | 2.83 ± 0.07 |
| I(0)* ($cm^{-1}$) from Guinier | 24.86 ± 0.036 | 85.5 ± 0.12 | 44.28 ± 0.056 | 60.24 ± 0.16 | 171.59 ± 0.15 | 8.66 ± 0.075 | 20.71 ± 0.044 |
| $R_g$ (nm) from P(r) | 2.096 ± 0.0003 | 2.88 ± 0.0005 | 2.62 ± 0.0006 | 4.34 ± 0.0014 | 2.95 ± 0.0003 | 2.70 ± 0.03 | 2.9 ± 0.05 |
| $D_{max}$ (nm) | 6.75 ± 0.67 | 9.6 ± 0.95 | 8.3 ± 0.83 | 15.5 ± 1.2 | 10 ± 1.0 | 8.5 ± 0.86 | 10.3 ± 1.2 |
| Porod volume ($nm^3$) | 52.4 | 65.9 | 72.44 | 114.07 | 66.47 | 62.53 | 60.82 |
| Dry volume calculated from sequence ($nm^3$)** | 38.468 | 55.077 | 63.126 | 92.784 | 55.059 | 55.89 | 55.076 |
| **Molecular mass determination (kDa)** | | | | | | | |
| From Porod volume ($V_p$/~1.6) | 24.9 | 47.6 | 40.9 | 76.1 | 48.5 | 39.1 | 42.7 |
| From SAXS MoW2*** | 28.9 | 49.7 | 35.7 | 65.2 | 50.9 | 44.6 | 46.4 |
| Bayesian inference | 28.9 | 46.6 | 41.9 | 67.1 | 46.6 | 42.8 | 40.2 |
| From I(0) using $V_c$ invariant | 28.2 | 43.5 | 43.4 | 65.6 | 43.7 | 40.2 | 42.7 |
| Calculated from sequence**** | 31.8 | 45.5 | 52.2 | 76.5 | 45.5 | 45.5 | 45.5 |
| Number of residues | 274 | 396 | 452 | 664 | 396 | 396 | 396 |
| **Software list** | | | | | | | |
| Primary data reduction | Automated pipeline at beamline | | | | | | |
| Data processing | PRIMUS (ATSAS v.2.8.0) | | | | | | |
| Ab initio analysis | DAMMIN and GASBOR | | | | | | |
| Fitting | CRYSOL | | | | | | |
| Model refinement | SREFLEX | | | | | | |
| Flexibility analysis | EOM 2.0 | | | | | | |
| Model superimpositions | SASpy plugin for Pymol | | | | | | |
| 3D graphics images | Pymol (v.1.8.2.1) | | | | | | |

*I(0) values shown in SEC-SAXS measurements vary depending on protein concentration at the analyzed peak, and are therefore not normalized to protein concentration. The structural parameters analyzed are independent of this value (i.e., Rg, $D_{max}$, volumes).
**http://biotools.nubic.northwestern.edu/proteincalc.html
***SAXS MoW2
****http://web.expasy.org/

Table S3, related to Figure 4. X-ray data.

| Crystal structure | 6HTF (rF10-SH2) |
|---|---|
| **Data collection** | |
| Space group | P 21 21 2 |
| Cell dimensions | |
| *a*, *b*, *c* (Å) | 145.53, 32.95, 80.63 |
| α, β, γ (°) (°) | 90, 90, 90 |
| Resolution (Å) | 50 (2.1) * |
| $R_{meas}$ | 10.2 (81.1) |
| $I / \sigma I$ | 12.99 (1.95) |
| Completeness (%) | 93.93 (84.43) |
| Redundancy | 3.89 (3.90) |
| **Refinement** | |
| Resolution (Å) | 2.1 |
| No. reflections | 22064 |
| $R_{work}$ / $R_{free}$ | 0.213 / 0.252 |
| No. atoms | 3041 |
| Protein | 2915 |
| Ligand/ion | 0 |
| Water | 126 |
| Protein residues | 362 |
| *B*-factors | |
| Protein | 40.75 |
| Ligand/ion | N/A |
| Water | 36.85 |
| R.m.s. deviations | |
| Bond lengths (Å) | 0.023 |
| Bond angles (°) | 1.44 |
| Ramachandran analysis | |
| Favored regions | 95.53% |
| Allowed regions | 4.47% |
| Outliers | 0 |

*Values in parentheses are for the highest-resolution shell.

Table S4, related to Figure S4. SAXS parameters of Btk-rF10 complexes.

| **Data collection parameters** | | | | | |
|---|---|---|---|---|---|
| Instrument | MD29 beamline, ESRF Grenoble - France | | | | |
| Wavelength (Å) | 0.9919 | | | | |
| q-range ($nm^{-1}$) | 0.03563 - 5 | | | | |
| Exposure time (sec) | 5 (10 frames x 0.5 sec) | | | | |
| Temperature (K) | 290 | | | | |
| **Samples** | **rF10** | **SH2-rF10** | **SH2-KD-rF10** | **SH3-SH2-KD-rF10** | **Full-length-rF10** |
| Measurement mode | SEC-SAXS | batch | SEC-SAXS | batch | batch |
| Concentration range ($mg.ml^{-1}$) | 100 µl at 14 | 0.6 – 2.7 | 100µl at 20 | 0.4 – 3.6 | 0.4 – 2.9 |
| SASBDB identifier | N/A | N/A | N/A | N/A | N/A |
| **Structural parameters** | | | | | |
| $R_g$ (nm) from Guinier | 2.42 ± 0.2 | 2.6 ± 0.3 | 3.46 ± 0.12 | 3.55 ± 0.26 | 4.48 ± 0.38 |
| I(0)* ($cm^{-1}$) from Guinier | 37.12 ± 0.06 | 35.7 ± 0.18 | 87.82 ± 0.24 | 61.09 ± 0.19 | 79.46 ± 0.23 |
| $R_g$ (nm) from P(r) | 2.47 ± 0.006 | 2.6 ± 0.01 | 3.52 ± 0.007 | 3.6 ± 0.01 | 4.69 ± 0.02 |
| $D_{max}$ (nm) | 7.9 ± 0.8 | 8.2 ± 0.8 | 11.5 ± 1.2 | 12.5 ± 1.2 | 16.9 ± 1.7 |
| Porod volume ($nm^3$) | 50.85 | 67.4 | 100.6 | 116.46 | 156.04 |
| Dry volume calculated from sequence ($nm^3$)** | 37.581 | 53.569 | 92.638 | 100.685 | 130.345 |
| **Molecular mass determination (kDa)** | | | | | |
| From Porod volume ($V_p$/~1.6) | 24.8 | 37.9 | 70.4 | 81.2 | 100.8 |
| From SAXS MoW2*** | 27.1 | 41.9 | 79.5 | 86.2 | 116.2 |
| Bayesian inference | 28.2 | 37.7 | 67.1 | 74.3 | 94.2 |
| From I(0) using $V_c$ invariant | 27.6 | 39.3 | 63.5 | 74.6 | 89.4 |
| Calculated from sequence**** | 31.1 | 44.3 | 76.6 | 83.2 | 107.7 |
| Number of residues | 274 | 391 | 670 | 726 | 938 |
| **Software list** | | | | | |
| Primary data reduction | Automated pipeline at beamline | | | | |
| Data processing | PRIMUS (ATSAS v.2.8.0) | | | | |
| Ab initio analysis | DAMMIN and GASBOR | | | | |
| Fitting | CRYSOL | | | | |
| Model refinement | SREFLEX | | | | |
| Flexibility analysis | EOM 2.0 | | | | |
| Model superimpositions | SASpy plugin for Pymol | | | | |
| 3D graphics images | Pymol (v.1.8.2.1) | | | | |

*I(0) values shown in SEC-SAXS measurements vary depending on protein concentration at the analyzed peak, and are therefore not normalized to protein concentration. The structural parameters analyzed are independent of this value (i.e., Rg, $D_{max}$, volumes).
**http://biotools.nubic.northwestern.edu/proteincalc.html
***SAXS MoW2
****http://web.expasy.org/

Table S5, related to STAR Methods. Oligonucleotides used for mutagenesis.

| Primer name | Sequence 5' to 3' (forward and reverse) |
|---|---|
| Btk K296E | For: GCTGAGCAACTGCTAGAGCAAGAGGGGAAAG<br>Rev: CTTTCCCCTCTTGCTCTAGCAGTTGCTCAGC |
| Btk Y223F | For: GTGGCCCTTTTCGATTACATGCCAATG<br>Rev: CATTGGCATGTAATCGAAAAGGGCCAC |
| Btk E301K | For: GCAAGAGGGGAAAAAGGGAGGTTTCATTGTC<br>Rev: GACAATGAAACCTCCCTTTTTCCCCTCTTGC |
| Btk R307G | For: GGTTTCATTGTCGGCGACTCCAGCAAAGC<br>Rev: GCTTTGCTGGAGTCGCCGACAATGAAACC |
| Btk K311E | For: CAGAGACTCCAGCGAGGCTGGCAAATATACAG<br>Rev: CTGTATATTTGCCAGCCTCGCTGGAGTCTCTG |
| Btk Q341A | For: GTTGTGTGTTCCACACCTGCGAGCCAGTATTACCTGGC<br>Rev: GCCAGGTAATACTGGCTCGCAGGTGTGGAACACACAAC |
| Btk H364D | For: CATTAACTACCATCAGGACAACTCTGCAGGACTC<br>Rev: GAGTCCTGCAGAGTTGTCCTGATGGTAGTTAATG |
| Btk S371P | For: CTGCAGGACTCATACCCAGGCTCAAATATCCAG<br>Rev: CTGGATATTTGAGCCTGGGTATGAGTCCTGCAG |
| Btk R372G | For: CTCTGCAGGACTCATATCCGGCCTCAAATATCCAG<br>Rev: CTGGATATTTGAGGCCGGATATGAGTCCTGCAGAG |
| Btk K374N | For: CTCTGCAGGACTCATATCCAGGCTCAACTATCCAG<br>Rev: CTGGATAGTTGAGCCTGGATATGAGTCCTGCAGAG |
| Btk C481S | For: GGTAGTTCAGGAGGCTGCCATTGGCCATGTA<br>Rev: TACATGGCCAATGGCAGCCTCCTGAACTACC |
| Btk D521N | For: GTTCCTTCACCGAAACCTGGCAGCTCG<br>Rev: CGAGCTGCCAGGTTTCGGTGAAGGAAC |
| Btk Y551F | For: GGATGATGAATTCACAAGCTCAGTAG<br>Rev: CTACTGAGCTTGTGAATTCATCATCC |

### KEY RESOURCES TABLE

| REAGENT or RESOURCE | SOURCE | IDENTIFIER |
|---|---|---|
| **Antibodies** | | |
| Mouse monoclonal anti-Total pY (clone 4G10) | Millipore | Cat# 05-321; RRID: AB_309678 |
| Mouse monoclonal anti-Btk (D6T2C) | Cell Signaling | Cat# 56044; RRID: AB_2799503 |
| Rabbit polyclonal anti-Btk | Thermo Scientific | Cat# PA5-27392; RRID: AB_2544868 |
| Mouse monoclonal anti-Btk (pY551)/Itk (pY511) Clone 24a | BD Biosciences | Cat# 558034; RRID: AB_2067823 |
| Rabbit polyclonal anti-Btk (pY223) | Cell Signaling | Cat# 5082; RRID: AB_10561017 |
| Rabbit polyclonal anti-p44/42 MAPK (Erk1/2) | Cell Signaling | Cat# 9102; RRID: AB_330744 |
| Mouse monoclonal anti-phospho-p44/42 MAPK (Erk1/2) (Thr202/Tyr204) (E10) | Cell Signaling | Cat# 9106; RRID: AB_331768 |
| Rabbit polyclonal anti-PLCγ2 | Cell Signaling | Cat# 3872; RRID: AB_2299586 |
| Rabbit polyclonal anti-PLCγ2 (Tyr1217) | Cell Signaling | Cat# 3871; RRID: AB_2299548 |
| Mouse monoclonal anti-Flag | Sigma | Cat# F3165; RRID: AB_259529 |
| Mouse monoclonal anti-Penta-his | Qiagen | Cat# 34660; RRID: AB_2619735 |
| Rabbit polyclonal anti-PLCγ2 (Tyr753) [EPR5914-3] | Abcam | Cat# ab133455; RRID: AB_2163712 |
| Mouse monoclonal anti-Tubulin | Sigma | Cat# T9026; RRID: AB_477593 |
| Mouse monoclonal anti-Myc-tag Myc.A7 DyLight800 | Thermo Scientific | Cat# MA1-21316-D800; RRID: AB_2536996 |
| Goat polyclonal anti-mouse IgG IRDye 800CW | LiCor | Cat# 926-32210; RRID: AB_621842 |
| Donkey polyclonal anti-Rabbit IgG (H+L) IRDye800 | Rockland | Cat# 611-732-127; RRID: AB_220158 |
| Goat polyclonal anti-Mouse IgG (H+L) Peroxidase AffiniPure | Jackson ImmunoResearch | Cat# 115-035-003; RRID: AB_10015289 |
| Goat polyclonal anti-Rabbit IgG (H+L) Peroxidase AffiniPure | Jackson ImmunoResearch | Cat# 111-035-003; RRID: AB_2313567 |
| **Chemicals, Peptides, and Recombinant Proteins** | | |
| Peptide FITC-ADNDpYIIPLPD | Eurogentec | N/A |
| Peptide Biotin-ERDINSLYDVSR-amide | Eurogentec | N/A |
| Ibrutinib (PCI-32765) | Selleck Chemicals | Cat# S2680 |
| AffiniPure F(ab') fragment goat anti-Human IgM, $Fc_{5\mu}$ fragment specific | Jackson ImmunoResearch | Cat# 109-006-129; RRID: AB_2337553 |
| AffiniPure F(ab') fragment goat anti-Human IgG, Fcγ fragment specific | Jackson ImmunoResearch | Cat# 109-006-098; RRID: AB_2337551 |
| Benzonase, *E.coli* recombinant | Biovision | Cat# 7680-5 |
| Cy 5 Annexin V | BD Biosciences | Cat# 559934 |
| 7-Amino-Actinomycin (7-AAD) | BD Biosciences | Cat# 559925 |
| **Critical Commercial Assays** | | |
| PureLink HiPure Plasmid Midiprep Kit | Invitrogen | Cat# K210004 |
| CellTiter-Glo 2.0 | Promega | Cat# G9241 |
| PolyFect Transfection Reagent | Qiagen | Cat# 301105 |
| FuGENE HD Transfection Reagent | Promega | Cat# E2311 |
| Quikchange II Site-Directed Mutagenesis Kit | Agilent | Cat# 200523 |
| **Deposited Data** | | |
| rF10-SH2 structure | This paper | PDB: 6HTF |
| SAXS data for human Btk Full-length | This paper | SASDF83 |
| SAXS data for human Btk SH3-SH2-KD | This paper | SASDF73 |
| SAXS data for human Btk SH2-KD | This paper | SASDF63 |
| SAXS data for human Btk KD | This paper | SASDF53 |
| **Experimental Models: Cell Lines** | | |

| | | |
|---|---|---|
| HEK293 | ATCC | Cat# CRL-1573; RRID: CVCL_0045 |
| HEK293T | ATCC | Cat# CRL-3216; RRID: CVCL_0063 |
| Sf9 insect cells | Thermo Scientific | Cat# 11496-015 |
| HBL-1 | Gift from M. Thome-Miazza | RRID: CVCL_4213 |
| DOHH2 | Gift from M. Thome-Miazza | Cat# ACC-47; RRID: CVCL_1179 |
| TMD8 | Gift from M. Thome-Miazza | RRID: CVCL_A442 |
| Experimental Models: Organisms/Strains | | |
| E.coli BL21 (DE3) | Thermo Scientific | Cat# EC0114 |
| E.coli Origami B (DE3) | Merck | Cat# 70837-3 |
| E.coli DH10B | Invitrogen | Cat# EC0113 |
| Oligonucleotides | | |
| Primers for site-derived mutagenesis, see Table S5 | This paper | N/A |
| Recombinant DNA | | |
| Plasmid pCS2-6xmyc | RZPD | N/A |
| Plasmid pETM30-6xhis-GST-TEV | EMBL | N/A |
| Plasmid pET21a | Novagen | Cat# 69740-3 |
| pFast-DUAL vector | Thermo Scientific | Cat# 10712024 |
| pFast-1 vector | Thermo Scientific | Cat# 10360014 |
| Plasmid pCW57.1 | Addgene | Cat# 41393 |
| Plasmid pMD2G | Addgene | Cat#12259 |
| Plasmid pCMVR8.74 | Addgene | Cat# 22036 |
| Plasmid pcDNA3.1 | Thermo Scientific | Cat# V79020 |
| Software and Algorithms | | |
| Image Studio Lite v5.2.5 | LiCor | www.licor.com |
| Prism 7 | GraphPad | www.graphpad.com |
| Geneious v9.1.8 | Biomatters | www.geneious.com |
| MassLynx 4.1 | Waters | www.waters.com |
| ATSAS Suite v2.8.3 | EMBL Hamburg | www.embl-hamburg.de/biosaxs |
| Ensemble Optimisation Method (EOM 2.0) | EMBL Hamburg | www.embl-hamburg.de/biosaxs |
| SASpy v2.8.0 | EMBL | www.embl-hamburg.de/biosaxs |
| MacPyMOL v1.8.2.2 | DeLano Scientific | www.pymol.org |
| BiKi LifeSciences suite 1.3 | Omics | www.omictools.com |
| Maestro (release 2016-1) | Schrodinger | www.schrodinger.com |
